# Synaptic circuits for irradiance coding by intrinsically photosensitive retinal ganglion cells

**DOI:** 10.1101/442954

**Authors:** Shai Sabbah, Carin Papendorp, Elizabeth Koplas, Marjo Beltoja, Cameron Etebari, Ali Noel Gunesch, Luis Carrete, Min Tae Kim, Gabrielle Manoff, Ananya Bhatia-Lin, Tiffany Zhao, Daniel Schreck, Henry Dowling, Kevin L. Briggman, David M. Berson

## Abstract

We have explored the synaptic networks responsible for the unique capacity of intrinsically photosensitive retinal ganglion cells (ipRGCs) to encode overall light intensity. This luminance signal is crucial for circadian, pupillary and related reflexive responses light. By combined glutamate-sensor imaging and patch recording of postsynaptic RGCs, we show that the capacity for intensity-encoding is widespread among cone bipolar types, including OFF types.

Nonetheless, the bipolar cells that drive ipRGCs appear to carry the strongest luminance signal. By serial electron microscopic reconstruction, we show that Type 6 ON cone bipolar cells are the dominant source of such input, with more modest input from Types 7, 8 and 9 and virtually none from Types 5i, 5o, 5t or rod bipolar cells. In conventional RGCs, the excitatory drive from bipolar cells is high-pass temporally filtered more than it is in ipRGCs. Amacrine-to-bipolar cell feedback seems to contribute surprisingly little to this filtering, implicating mostly postsynaptic mechanisms. Most ipRGCs sample from all bipolar terminals costratifying with their dendrites, but M1 cells avoid all OFF bipolar input and accept only ectopic ribbon synapses from ON cone bipolar axonal shafts. These are remarkable monad synapses, equipped with as many as a dozen ribbons and only one postsynaptic process.

## Introduction

Environmental light intensity (irradiance) synchronizes the circadian clock, constricts the pupil, and regulates hormones and mood^1,2^. Most retinal ganglion cells (RGCs) do not encode intensity faithfully. Light steps affect their activity only briefly and their maintained firing rates are uncorrelated with irradiance^3^. However, intrinsically photosensitive retinal ganglion cells (ipRGCs) do effectively encode irradiance. They are ON cells with maintained firing rates that remain proportional to light intensity essentially indefinitely^4^. Like other RGCs, ipRGCs receive photoreceptor signals through synaptic input from bipolar cells. However, they also sense light directly, through melanopsin phototransduction^4-9^. Both excitatory influences encode luminance. This implies that bipolar cells driving ipRGCs encode intensity, but it is unknown which bipolar types are involved and whether they are uniquely suitable for this purpose.

Each type of bipolar cell (BC) distributes its axonal output to a different level of the inner plexiform layer (IPL) of the retina^10,11^, and makes excitatory glutamatergic ribbon contacts with distinct sets of RGCs and amacrine cells (ACs)^12^. Some BCs are excited by light increments (ON-BCs), other by decrements (OFF-BCs)^13-15^. Some draw input mainly from rods, others from cones^16^. Light adaptation in photoreceptors enhances temporal contrast^17^ and further temporal filtering occurs in bipolar cells. But while some BC types are more sustained than others^14,18,19^, it is unclear whether any are specialized to encode absolute light intensity and whether they contact ipRGCs selectively.

Dendritic stratification of ipRGCs provides some clues. The five ipRGCs types (M1 through M5), which all exhibit sustained ON responses^7^, deploy their dendrites in one or both of two IPL sublaminae^20^. One lies within the inner tier of the ON sublayer, where relatively sustained ON bipolar cells terminate^12,21,22^. The other, surprisingly, occupies the distal margin of the OFF sublayer. The ipRGCs that stratify there (M1 and M3 cells) are also ON cells. They apparently receive ectopic ribbon contacts from ON bipolar axons as they descend toward their main axonal terminal arbor in the ON-IPL^23,24^. Some of these are atypical bipolar synapses, with only a single postsynaptic partner at the ribbon synapse (“monad”)^25^ whereas in the classical “dyad” synapse there are two. Typically, one postsynaptic process at dyad synapses is an amacrine cell^26^, which can contribute to temporal filtering by inhibitory feedback. The form, targets, and functional properties of ON bipolar output synapses in this ‘accessory ON sublayer’ remain largely mysterious.

Here, we characterize synaptic structure and intensity encoding within the BC-to-ipRGC circuit. By electron microscopic reconstruction, patch-clamp recording, and glutamate imaging, we characterize the bipolar input to ipRGC dendrites in both IPL layers. In the accessory ON sublayer, large, monad multi-ribbon synaptic plaques from Type 6 cone BCs predominate; Types 7-9 also provide some monad inputs. These same bipolar types (6-9) contact ipRGCs dendrites in the conventional ON sublayer, but at dyad synapses; Type 6 again dominates. Specialized bipolar inputs cannot fully account for why most RGCs encode intensity poorly because we find sustained and intensity-dependent glutamate release from all cone bipolar cells throughout all IPL layers, both ON and OFF. The intensity-encoding capacity of ipRGCs thus arises partly because they lack high-pass postsynaptic filtering that other RGCs perform on this bipolar input.

## Results

ipRGCs were reported to receive sustained and intensity-encoding excitatory synaptic input ^27^. However, a systematic characterization of the excitatory synaptic input onto each of the ipRGC types has never been conducted. Here, recording excitatory currents in whole-cell configuration, we show that all ipRGCs receive excitatory sustained and intensity-encoding synaptic input, regardless of their synapse type with BCs or whether they stratify in the ON, OFF, of both IPL sublamina, (Extended Data Figure 1).

### Ectopic synapses appear predominantly in Type 6 bipolar cells

We used serial block-face electron microscopy (SBEM) to determine the synaptic drive to ipRGCs. We utilized a SBEM volume of adult mouse retina that extended through the IPL, and was stained to reveal organelles^28^, allowing us to identify synaptic vesicles and ribbons. We reconstructed hundreds bipolar cells (BCs) in the volume. Extended Data Figure 2 shows all reconstructed BCs per type, along with their stratification patterns and mosaics. In addition to the 13 known cone BC types and a single rod BC type^11,29^, we identified another cone BC type – Type 0 ^30^. Figure 1a-c shows representatives examples of each BC type and group data on their terminal field area and IPL stratification depth. By scanning the far outer IPL, we identified over 500 ectopic ON cone bipolar ribbon synapses (Fig. 1d-f). These were almost exclusively monads (Extended Data Figure 3a). In each synapse, we located multiple ribbons (gray stripes), and each ribbon was surrounded by vesicles (black aggregates). See Extended Data Figure 3 for additional examples of ectopic synapses identified on BC Types 6, 7, 8, and 9. Type 6 BCs were the predominant source of ectopic synapses (Fig. 1e,g).

**Figure 1.**
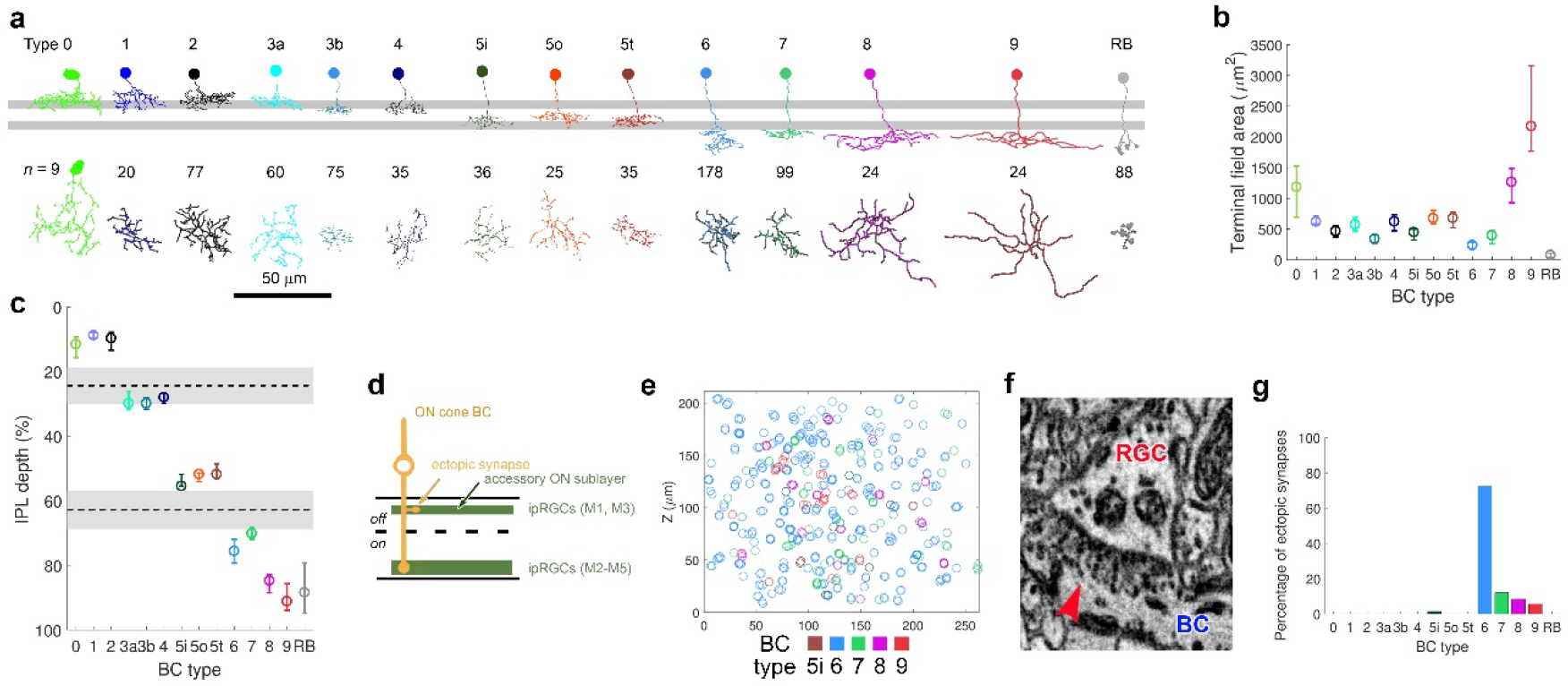
Ectopic synaptic input is ubiquitous and unique to ipRGC networks. **a**, Representatives of the different BC types encountered in the SBEM volume, encompassing the complete BC repertoire in mice plus the new Type 0 BC. Two horizontal gray stripes mark the ON and OFF cholinergic bands, as estimated from the ON and OFF dendritic arbors of ON-OFF-DSGCs (*n* = 15). **b**,**c**, Different BC types exhibit characteristic terminal field area (**b**) and stratification depth across the IPL (**c**). Dashed lines and shaded gray areas mark the median and first and third quartiles of the cholinergic bands’ IPL depth. **d**, Schematic of ectopic ribbon synapse from a BC onto the dendrites of ipRGCs. Ectopic synapses can be found at the INL-IPL boundary (IPL depth 5%-25%). **e**, Distribution of ectopic synapses in the SBEM volume as a function of the BC type that gives rise to them. **f**. Example electron micrograph of an ectopic synapse on the shaft of a type 6 BC onto a RGC’s dendrite. Seven ribbons (gray stripes) surrounded by vesicles (black clamps) can be seen. **g**, Type 6 BCs are the predominant source of ectopic synapses.

### BC Types 6 and 7 provide the majority of input to ipRGCs through ectopic monad and conventional dyad synapses

We identified in the volume all the ipRGC types, except for the rare M3 cells^31^. To support our identification, we compared key morphological statistics of SBEM traces to those of genetically, morphologically, and physiologically identified ipRGCs (Extended Data Figure 4). Figure 2a shows an example of an M1 cell (black) that stratifies in the OFF-IPL, and therefore, is in position to receive ectopic synapses. Also depicted are the reconstructions of bipolar cells that make ectopic synapses with it (different types depicted in different colors), and the location of ribbon synapses (red dots). Type 6 BCs were the source of over 80% of all ectopic inputs onto M1 cells (Fig. 2e). In fact, all bipolar input to M1 cells came from ectopic synapses; they actively avoided axon terminals from OFF-BCs in the OFF-IPL, and there were no apparent ribbon synaptic contacts at all onto the proximal dendrites of M1 cells within the ON sublamina of the IPL. M2, M4, and M5 ipRGCs, stratifying only in the ON-IPL, received their cone bipolar input through terminal dyad synapses, mainly from the axon terminals of BC Types 6 and 7 (Fig. 2b-d,f-h). Overall, it appears that bipolar inputs to ipRGCs derive predominantly from cone BC Types 6 and 7, but that these inputs have a unique form (multiribbon monads) and location (accessory ON sublayer) for M1 cells than for other ipRGC types.

**Figure 2.**
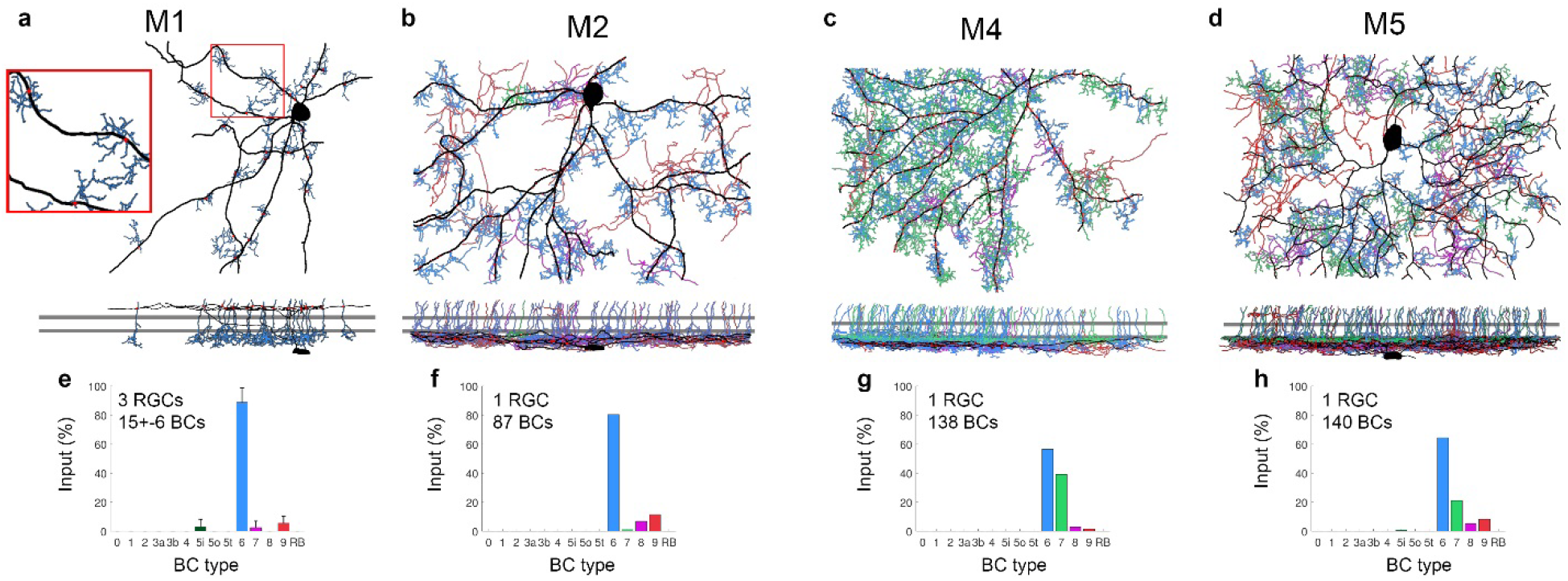
Cone BC Types 6 and 7 provide the bulk input to ipRGCs through either ectopic or conventional synapses. **a-d**, SBEM reconstruction of BC input onto M1, M2, M4, and M5 ipRGCs, en face view (top) and side view (bottom). RGCs are depicted in black; each BC type is depicted in a unique color. Ribbon synapses are depicted as red circles. Two horizontal gray stripes mark the ON and OFF cholinergic bands. **e**-**h**, Percent input of each BC type to the ipRGC in question.

Of note, ectopic synaptic input was also identified on a previously unknown bistratified RGC type. This RGC received ectopic input predominantly from BC Type 6, but also terminal dyad synapses from BC Types 1, 2, 6, 8, and 9 (Extended Data Figure 5). Ectopic synaptic input was also identified onto several mono- and bistratified AC types. We did not characterize further the synaptic input to these ACs.

The relationship between the densities of BCs to those of synapses differed across ipRGC types. M1 ipRGCs showed a 1:1 ratio between number of presynaptic BCs and number of input synapses. However, in all other ipRGC types, the number of synapses was ~1.6 greater than that of presynaptic BCs. Each BC typically made one or two synapses on a given ipRGC, though we saw as many as four synapses in a few cases. These synapse and BC densities qualitatively correlated with the amplitude of steady-state light-evoked excitatory currents in ipRGCs (Extended Data Figure 6).

The preceding results demonstrate that ipRGCs, which are uniquely light intensity-encoding, are also distinctive in their bipolar input (mostly BC Types 6 and 7), raising the question of whether these bipolar types are uniquely specialized to handle intensity information. Knowing the stratification depth of these BC types, we next studied the glutamate release at the stratification depths of BC Types 6 and 7 and compared it to glutamate release from other BCs across the IPL.

### Sustained intensity-encoding glutamate release at light onset across IPL

We measured the glutamate released from BCs (and ACs) at 10 different depths within the IPL, from the inner nuclear layer (INL; IPL depth 0%) to the ganglion cell layer (GCL; IPL depth 100%). To measure the glutamate level, we used iGluSnFR, a membrane-targeted genetically-encoded indicator for glutamate that is extremely sensitive and fast ^32^. Extended Data Figure 7 shows a two-photon z-stack of the iGluSnFR-expressing dendrites of RGCs and ACs in each examined IPL depth. Glutamate release was measured in response to 30 sec of light, and at 5 different intensities. Based on our SBEM data, we could tell which BC types stratify at each IPL depth. For example, Types 6 and 7 that predominate the input to ipRGCs stratify in 65-75% IPL depth. As expected, glutamate release at these IPL depths persisted for the entire 30-sec stimulus duration. The steady-state amplitude, estimated as the average response over the last five seconds of the stimulus, correlated with light intensity (Fig. 3a, IPL depths 65% and 75%). However, in a major surprise, we found that all BCs encode intensity in their synaptic output (Fig. 3a). The glutamate signal increased monotonically in the ON-IPL with increasing stimulus intensity, but showed a monotonic decrease in the OFF-IPL (Fig. 3a).

**Figure 3.**
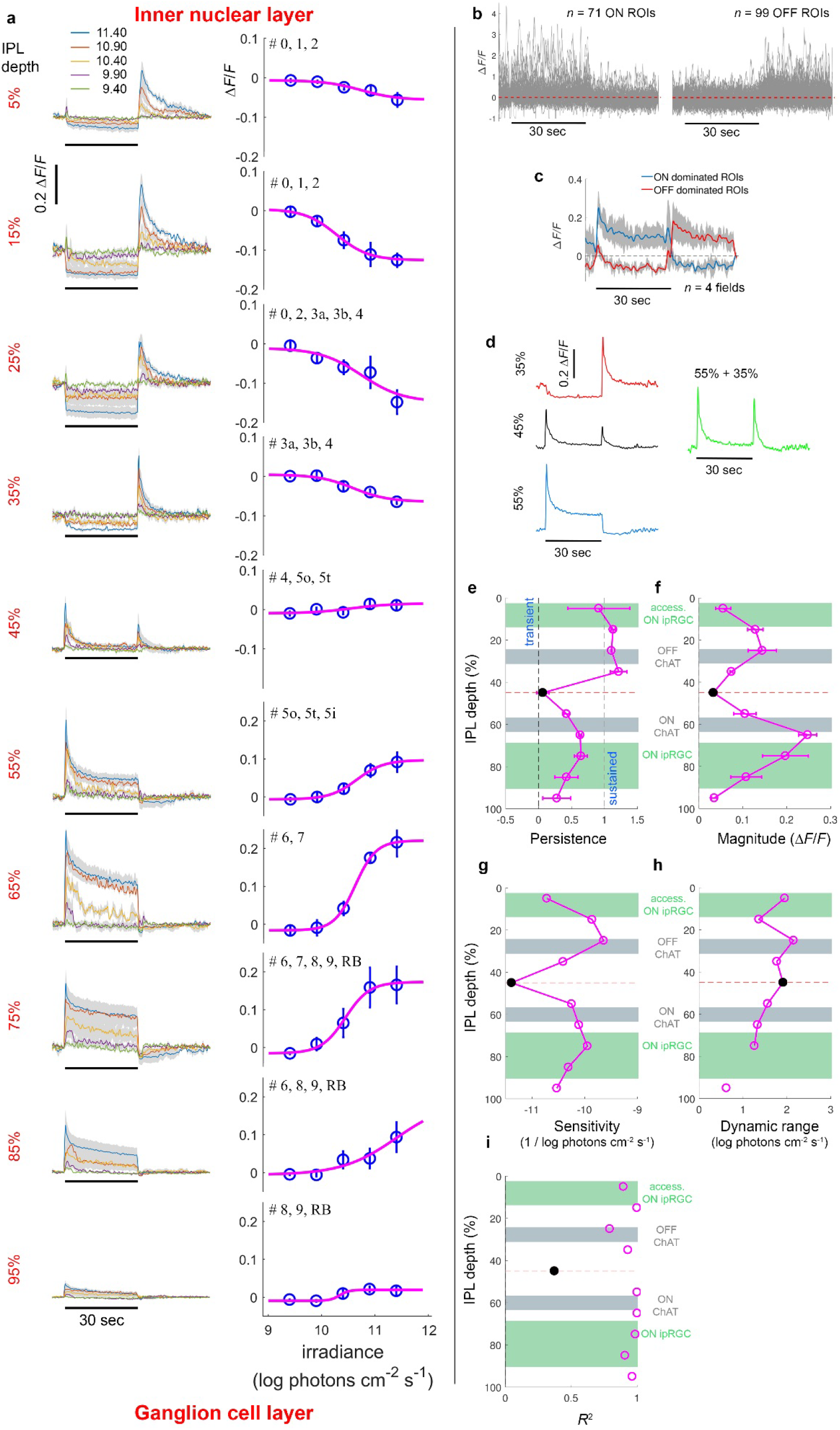
Glutamate release across the IPL is sustained and intensity-encoding. **a**. Glutamate release from BCs and ACs at 10 different depths across the IPL, from the inner nuclear layer (IPL depth 0%, top) to the ganglion cell layer (IPL depth 100%, bottom). Glutamate release was quantified using the fluorescence indicator iGluSnFR, and measured in response to 30 sec of light (indicated by a horizontal black bar) and at 5 different light levels (log photons cm^-2^ s^-1^; depicted in different colors). Left column shows the glutamate response (Δ*F*/*F*) as a function of time for five light stimulus intensities, for each IPL depth. Right column shows the steady-state glutamate response (Δ*F*/*F*) as a function stimulus intensity, for each IPL depth. Steady-state response was estimated as the mean response over the 5 last seconds of the light phase of the stimulus. BC types that stratify at each IPL depth (whose mean normalized density exceeded 20%) are indicated in black. In the inner IPL (55%-95% IPL depth), the glutamate response typically started with a large-amplitude transient, and then gradually rolled off until the termination of the stimulus, upon which glutamate release abruptly dropped to, or at certain IPL depths, below baseline. Additionally, at certain inner IPL depths, the glutamate release became more sustained as the stimulus intensity increased. In contrast, in the outer IPL (5%-35% IPL depth), glutamate release dropped abruptly at the onset of the stimulus, and stayed virtually constant until the termination of the stimulus, upon which glutamate release abruptly increased above baseline and spiked. **b**,**c**. The glutamate release, analyzed separately for selected identified single dendrites, peaked either at light onset or offset, but never at both. **d**. Summing the glutamate release at the two IPL depths (35% and 55%) spanning inner-outer IPL transition depth (45% IPL depth) produced a trace similar to that observed at the 45% transition depth. **e**. Glutamate release in the outer IPL is almost perfectly sustained. However, glutamate release in the inner IPL is most sustained at the middle of the inner IPL (IPL depths 65% and 75%), and becomes more transient toward the ON-OFF IPL boundary and the IPL-GCL boundary. **f**. The magnitude of steady state glutamate release (calculated as the maximum – minimum glutamate release encountered across the stimulus intensities) was minimal at the ON-OFF IPL boundary, largest at the middle of the outer and inner IPL (IPL depths 25% and 65%, respectively), and decreased toward both IPL edges. The depths of highest magnitude coincide with the ON and OFF ChAT bands. **g**. Sensitivity of glutamate release (see Methods) was minimal at the ON-OFF IPL boundary, largest at the middle of the outer and inner IPL, and decreased toward both IPL edges. **h**. The dynamic range of steady state glutamate responses (see Methods) varied slightly across the IPL. **i**. Goodness of fit to the Naka-Rushton function, was high across the IPL, except for the ON-OFF transition depth.

The only exception to this trend was observed near the middle of the IPL, at the transition between the ON and OFF sublayers (IPL depth 45%). Glutamate release at this ON/OFF transition zone exhibited two peaks, one at the onset and the other at the offset of light, and little if any steady state response (Fig. 3a, middle left). However, we suspected that at this depth, the global iGluSnFR signal, summed over the whole field of view, might poorly reflect the behavior of individual BCs, because it accounts for the sum of contributions of the closely interspersed ON and OFF bipolar signals. Indeed, the glutamate release, analyzed separately for selected identified single dendritic segments, peaked either at light onset or offset, but never at both (Fig. 3b,c). Moreover, we found that summing the glutamate indicator signals at the two IPL depths closest to the ON/OFF transition zone but exhibiting pure OFF or pure ON responses (35% and 55% depths, respectively) resulted in a trace similar to that observed at the ON/OFF transition zone (45% depth, Fig. 3d). Therefore, the glutamate release measured at the ON/OFF transition zone likely represents the summed contributions of ON and OFF bipolar signals. In the outer OFF sublayer (depths 5%-25%), we observed a transient glutamate release at the light onset. These responses might reflect the glutamate release from ON-BCs through ectopic synapses.

To summarize how the capacity for intensity encoding varies across the IPL, we developed two indices. First, we calculated the persistence index, which accounts for the ratio between the end and beginning of the response (see methods). Glutamate release in the OFF-IPL was almost perfectly sustained. However, glutamate release in the ON-IPL was most sustained at the middle of the inner IPL, corresponding only modestly with the stratification depth of ipRGCs (Fig. 3e). We also estimated the magnitude of the steady state glutamate release by calculating the difference in glutamate release encountered across the stimulus intensities. This magnitude was largest at the middle of the OFF and ON-IPL, again, not matched entirely to the stratification depths of ipRGCs (Fig. 3f). Fig. 3g-i presents the sensitivity and dynamic range of the steady-state response as well as the goodness of the Naka-Rushton fit. Taken together, these results suggest that differences between bipolar types that stratify at different IPL depths cannot account completely for the differences in intensity encoding between ipRGCs and conventional RGCs. This points to postsynaptic mechanisms as the primary basis of the filtering, which will be studied further below.

### Effect of inhibition on glutamate release

We repeated this experiment while blocking GABAergic and glycinergic amacrine inhibition using picrotoxin and strychnine, effectively revealing the output of BCs unfiltered by amacrine-cell influence. The effect of feedback inhibition on the persistence and capacity for intensity-encoding was minimal, if any (Extended Data Fig. 8), suggesting that these processes are shaped primarily by glutamatergic signaling in the outer retina and intrinsic membrane properties and synaptic release mechanisms at the BC axon terminals.

### Sustained glutamate release at light offset across IPL

To study the glutamate released from OFF-BCs when presented with their preferred contrast, we measured again the glutamate release across the IPL, but now with dark spots of different intensities over a fixed, bright background (Extended Data Fig. 9). As expected, the onset of the dark spots increased with glutamate release, and the later persisted throughout the 30-sec duration of the stimulus. However, we were unable to reproduce the evidence from Fig. 3 (bright spots) that OFF-BC output is intensity encoding, likely due to practical reasons. Specifically, technical limitations of our experimental setup and the need to limit the exposure of the wholemount retinal preparation to extended periods of light stimulation (a measurement profile across the IPL lasted ~3 hours), limited the range of stimulus intensities we could use.

### Intensity information in bipolar signals is filtered out by multiple postsynaptic mechanisms in conventional RGCs

Our results suggest that postsynaptic mechanisms are the primary basis of the filtering of intensity signals on their way to conventional RGCs. To explore the specific transmission stages involved in such filtering, we measured the glutamate release **onto** identified RGC types, and simultaneously, we also measured their postsynaptic excitatory currents and firing rate. We repeated these experiments for two RGCs, an ipRGC and a conventional RGC. The first type is the M4 ipRGC which stratifies at the ON plexus of ipRGCs (Fig. 4a,e). To measure the glutamate released onto M4 cells, we utilized the Opn4^Cre/+^ mouse that selectively labels ipRGCs^20^. Intravitreal injections of Cre-dependent iGluSnFR AAVs allowed targeting of these cells. We patched an iGluSnFR-positive cell with red dye in the pipette, and imaged regions of its arbor identified by their red dye fill. Because we had already patched the cell, we could simultaneously measure the excitatory currents and firing rate of the cell. The conventional RGC type examined is the ON direction selective ganglion cell (ON-DSGC) which stratifies with the ON cholinergic band and to a limited extent also with the OFF cholinergic band (Fig. 4e). ON-DSGCs are perhaps the most sustained among conventional RGCs. To measure the glutamate released onto ON-DSGCs, we utilized the Pcdh9^Cre/+^ mouse that selectively labels these cells (unpublished data; Brendan Lilley, Alex Kolodkin, D.M.B, S.S). Intravitreal injections of Cre-dependent iGluSnFR AAVs allowed targeting of ON-DSGCs (Fig. 4j); we confirmed their direction selectivity using drifting sinusoidal gratings stimuli (Fig. 4p) as well as morphology using depth series imaging.

**Figure 4.**
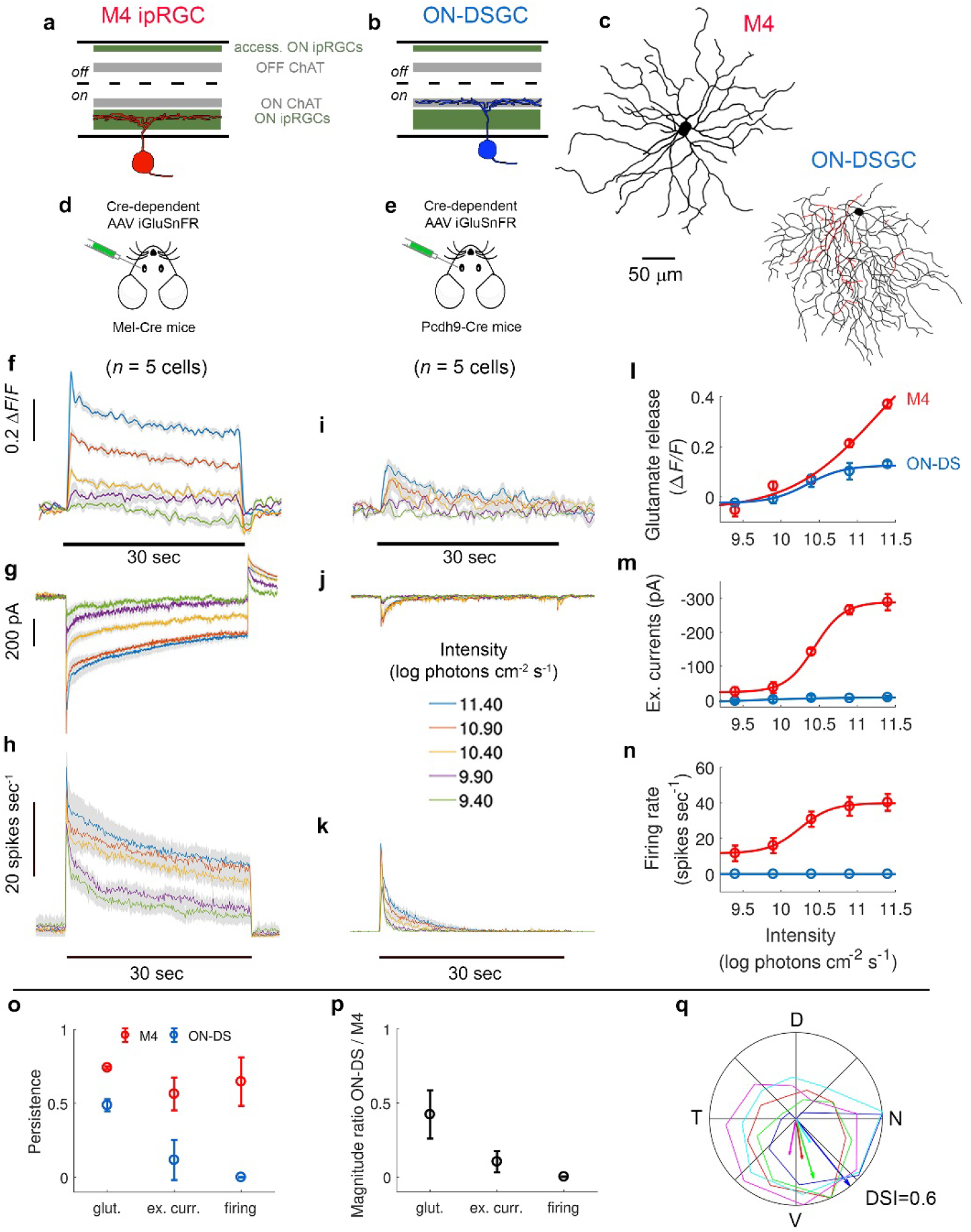
Capacity for intensity encoding decreases postsynaptically in ON-DSGCs more than in M4 ipRGCs. **a**,**b**, Two RGC types selected for further intensity-encoding characterization, M4 ipRGCs stratify at the ON plexus of ipRGCs (**a**) while ON-DSGCs stratify at the ON cholinergic band and to a limited extent also at the OFF cholinergic band (**b**). **c**, Exmaple somatodendritic reconstuctions of an M4 and ON-DSGC. **d**,**e**, Intravitreal injections of Cre-dependent iGluSnFR AAVs in either Mel-Cre or Pcdh9-Cre mice allowed targeting of M4 ipRGCs and ON-DSGCs, respectively. **f**-**h**, Glutamate release, excitatory currents, and firing rate for M4 ipRGCs in response to 30-sec bright spots over dark background, as a function of stimulus intensity. **i**-**j**, Glutamate release, excitatory currents, and firing rate for ON-DSGCs. **i**-**n**, Intensity-response curves for the glutamate release onto (**f**), and excitatory currents (**g**) and firing rate (**h**) of M4 ipRGCs (red) and ON-DSGCs (blue). **o**, Response persistence of M4 ipRGCs and ON-DSGCs. Response perseietence of ON-DSGCs was highest at the glutamate release, lower at the excitatory currents, and lowest at the firing rate. In contrast, response persistence on M4 ipRGCs decreased only slightly when moving between the signal transmission stages. **p**, Ratio of the steady-state magnitude between M4 ipRGCs and ON-DSGCs was higher for glutamate release, lower for excitatory currents, and lowest for firing rate. **q**, Polar plot shows response amplitude (normalized to maximum) for each direction; each cell depicted in a different color. The direction and length of vectors indicate the preferred direction and direction selectivity index (DSI) of cells, respectively (N, nasal; D, dorsal; T, temporal; V, ventral).

Glutamate release onto both cell types was sustained and increased with intensity (Fig. 4f,I,l). However, while the excitatory currents in M4 cells persisted for the entire 30-sec duration of the stimulus, those of ON-DSGCs were very transient, and quickly approached baseline (Fig. 4g,j,m). The firing rate showed similar trends (Fig. 4h,k,n). Figure 4g-h,i-k shows the glutamate release, excitatory currents and firing rate over time for the M4 ipRGCs and ON-DSGCs; Figure 4l-n compares the intensity-response curves of both cell types and across the three inspected levels of transmission. These results demonstrate that the intensity signal is being strongly filtered already at the level of excitatory currents.

This is reflected in two additional measures. First, the response persistence of ON-DSGCs, but not that of M4 ipRGCs, differed significantly and decreased abruptly when moving from glutamate release, through excitatory currents, to firing rate (Fig. 4o). Additionally, the ratio between the steady-state response magnitude of ON-DSGCs and M4 cells differed significantly across transmission stages and decreased sharply when moving from glutamate release, through excitatory currents, and to firing rate (Fig. 4p). This indicates that the effect of different stimulus intensities declined more for the ON-DSGCs when moving across the transformation stages. Therefore, the filtering of intensity signals involves multiple post-synaptic mechanisms, and is being executed at several transmission stages. Some of these mechanisms might include desensitization of glutamate receptors, variation in the spike generator, and feedforward inhibition from amacrine cells.

### Filtering of intensity signals *en route* to conventional OFF RGCs

ON-DSGCs are physiologically ON cells. They primarily, if not exclusively, respond at light onset. To test whether intensity signals on their way to OFF RGCs are also being strongly filtered already at the level of excitatory currents, we characterized the capacity for intensity-encoding for two additional conventional RGCs that stratify either at both ON and OFF ChAT bands (ON-OFF-DSGC) or inter-ChAT region (OFF *α*RGCs).

*ON-OFF direction selective ganglion cells (ON-OFF-DSGCs)*. ON-OFF-DSGCs costratify with the ON and OFF cholinergic bands. Reconstructing a similar ON-OFF-DSGC in our SBEM volume revealed that this cell receives input predominantly from the ON-BC Types 5o, 5t, and 5i as well as OFF-BC Types 3a, 3b, and 4 (Fig. 5u,v). As for ipRGCs, comparison of key morphological statistics of SBEM traces to those of genetically, morphologically, and physiologically identified RGCs supported our identification of conventional RGCs in the SBEM volume (Extended Data Figure 10). To target ON-OFF-DSGCs, we utilized intravitreal injections of non-specific iGluSnFR AAVs in wildtype mice. We patched a cell with red dye in the pipette, and imaged regions of its arbor that included dendrites that express iGluSnFR and are filled with dye. We confirmed their morphology based on 2-photon depth series as well as their direction selectivity using drifting sinusoidal gratings stimuli (Fig. 5z). We first optically recorded the glutamate release onto dendrites in the ON-IPL in response to light spots over a dark background. Glutamate release, but not excitatory currents, was sustained and intensity encoding (Fig. 5a-c). Next, we recorded the glutamate release onto dendrites in the OFF-IPL in response to light spots over a dark background. The withdrawal of glutamate release in response to light was sustained and intensity-encoding (Fig. 5d,e). Lastly, recording the glutamate release onto dendrites in the OFF-IPL, but this time, in response to dark spots over a bright background revealed that glutamate release increased at the onset of the dark spot, lasted for the entire stimulus duration, and correlated with the spot’s intensity (Fig. 5f-h). In summary, the glutamate released onto these ON-OFF-DSGCs was sustained and intensity-encoding, and either increased (light spots, ON arbor), decreased (light spots, OFF arbor), or increased (dark spots, OFF arbor). In contrast to the sustained and intensity-encoding glutamate release, excitatory currents transiently peaked at both the onset and offset of the stimulus, for either bright or dark spots. This reflects the summation of input from both ON and OFF dendritic arbors (Fig. 5b,g). *OFF α*RGCs. Next, we chose an OFF RGC type that stratifies between the ON and OFF ChAT bands (inter-ChAT, 35% IPL depth). Reconstructing a similar OFF *α*RGC in our SBEM volume revealed that this cell receives input from OFF-BC Types 3a, 3b and 4 (Fig. 5w,x). In a wildtype mouse intravitreouslly injected with non-specific iGluSnFR AAVs, we patched a cell with red dye in the pipette, and imaged regions of its arbor identified by their red dye fill. The glutamate released onto this OFF *α*RGCs in response to bright spots over a dark background mirrored our results when we averaged across all dendrites at the same IPL depth (compare Fig. 5i,k and Fig. 3 at 35% IPL depth). The reduction in glutamate was sustained and intensity-encoding. Additionally, recording the glutamate release onto the same OFF-IPL dendrites, but this time in response to dark spots over a bright background demonstrated that glutamate release increased at the onset of the dark spot, lasted for the entire stimulus duration, but did not show obvious correlation with the spot’s intensity (Fig. 5l,n). The excitatory currents showed similar but somewhat reduced trends, for both stimulus types (Fig. 5j,k,m,n).

**Figure 5.**
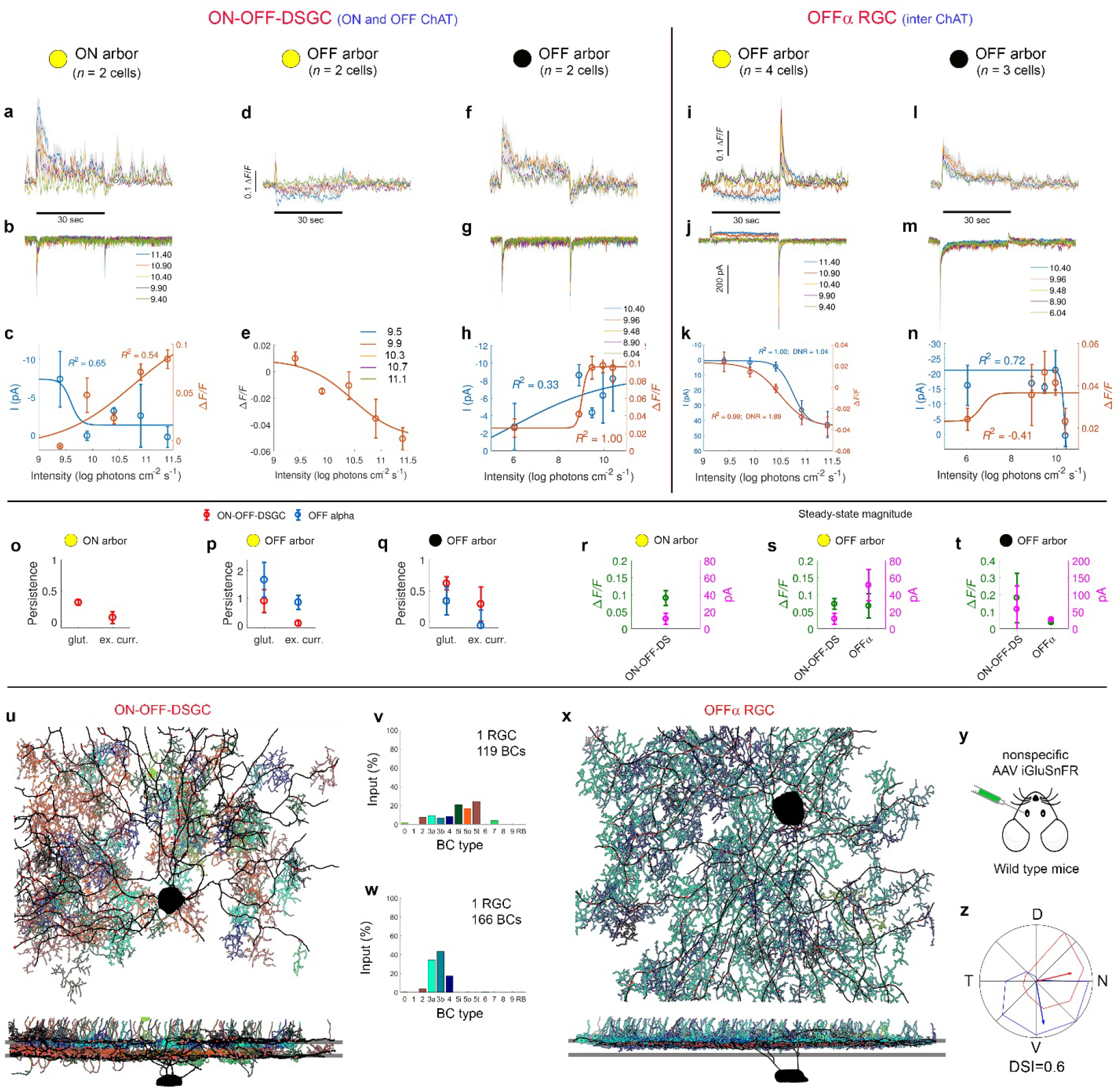
Sustained intensity-encoding glutamate release onto identified conventional RGCs. **a**-**h**, Intensity encoding capacity of ON-OFF-DSGCs as assessed using either bright spots on a dark background (**a**-**c**) or dark spots on a bright background (**d**-**h**). **a**-**b**, Simultaneous excitatory currents and glutamate release onto ON-IPL dendrites of ON-OFF-DSGCs in response to bright spots over a dark background. Glutamate release (Δ*F*/*F*) onto ON IPL dendrites of ON-DSGCs was measured in response to 30 sec of light (indicated by a horizontal black bar and specified in the legend) and at 5 different light levels (log photons cm^-2^ s^-1^; depicted in different colors). Excitatory currents were measured concurrently with the optical recording of glutamate, while holding the membrane voltage at the chloride reversal potential. **c**, Steady-state response as a function of stimulus intensity, for glutamate release (Δ*F*/*F*, orange) and excitatory currents (pA, blue, y-axis inverted to facilitate comparison to glutamate response). Steady-state response was estimated as the mean response over the 5 last seconds of the light phase of the stimulus. Goodness of fit to the Naka-Rushton function is indicated. **d,e**, Glutamate release onto OFF-IPL dendrites of ON-OFF-DSGCs; conventions similar to (**a**,**c**). **f**-**h**, Simultaneous excitatory currents and glutamate release onto OFF-IPL dendrites of ON-OFF-DSGCs in response to dark spots of five intensities (depicted by different colors) over a bright background (12.5 log photons cm^-2^ s^- 1^); conventions similar to (**a**-**c**). **i**-**n**, Intensity encoding capacity of OFFα RGCs in response to either bright spots on a dark background (**i**-**k**) or dark spots on a bright background (**l**-**n**). Conventions similar to (**a**-**c**). **o**-**q**, Persistance of glutamate release (glut.) and excitaotory currents (ex. curr.) in response to bright spots over a dark background (**o**,**p**) and dark spots over a bright background (**q**). Glutamate release readings are indicated for dendrites in the ON arbor (**o**) and OFF arbor (**p**,**q**). **r**-**s**, Magnitude of steady-state glutamate release (left y-axis, green) and excitaotory currents (right y-axis, magenta) in response to bright spots over a dark background (**r**,**s**) and dark spots over a bright background (**t**). Glutamate release readings are indicated for dendrites in the ON arbor (**r**) and OFF arbor (**s**,**t**). **u**, SBEM reconstruction of BC input onto an ON-OFF-DSGC, as viewed at the plane of the retina (top) and an orthogonal view (bottom). RGC is depicted in black; each BC type is depicted in a unique color. Ribbon synapses are depicted as red circles. Two horizontal gray stripes mark the ON and OFF cholinergic bands. **v**, Percent input of each BC type to the presumptive ON-OFF-DSGC in (**t**). **w**,**x**, SBEM reconstruction of BC input onto an OFFα RGC. **y**, Glutamate release onto RGCs was measured following intravitreal injections of non-specific iGluSnFR AAVs into the eyes of wildtype mice. Selected iGluSnFR-positive cells were patched with red dye in the pipette, and their arbor, identified by the red dye fill, was imaged. **z**, Direction selectivity of ON-OFF-DSGCs. Polar plot shows response amplitude (normalized to maximum) for each direction; each cell depicted in a different color. The direction and length of vectors indicate the preferred direction and direction selectivity index (DSI) of cells, respectively (N, nasal; D, dorsal; T, temporal; V, ventral).

Our results for the three conventional RGC types examined (ON-DSGC, ON-OFF-DSGC, and OFF *α*RGCs) indicate that the key locus for temporal filtering of sustained intensity-encoding signals is postsynaptic to the bipolar axon terminal. However, while the intensity signal in ON-DSGCs and ON-OFF-DSGCs is being strongly filtered as the level of excitatory currents, the signal in OFF *α*RGCs is largely retained in the excitatory currents. Thus, different conventional RGC types employ different strategies for filtering of intensity signals.

### Ectopic synapses transmit sustained and intensity-encoding signals

Synthesis of our connectomics and glutamate imaging findings for M4 ipRGCs suggests the capacity of intensity-encoding in BC types 6 and/or 7. M4 ipRGCs stratify in the ON-IPL (65%- 75% IPL depth), and therefore receive their input only from terminal dyad synapses. In contrast, M1 ipRGCs stratify in the far OFF-IPL (5-15% IPL depth) and thus receive their input only through ectopic monad synapses (Fig. 2a,e). We showed that excitatory currents in these M1 cells, that integrate rod/cone synaptic input with intrinsic melanopsin sensitivity, are sustained and intensity encoding (Extended Data Figure 1). However, the kinetics of glutamate release at ectopic synapses has never been examined directly.

To fill this gap, we recorded the glutamate released from ectopic synapses in the OFF-IPL. Our connectomics data show that these ectopic synapses relay input from ON-BCs to M1 ipRGCs and a novel bistratified RGC (Extended Data Figure 4). Such ectopic synapses might relay ON-BC input also to M3 ipRGCs that have not been identified in our SBEM volume. Intravitreal injections of Cre-dependent iGluSnFR AAVs in the Opn4^Cre/+^ mouse labeled only a limited number of dendrites at the far OFF-IPL, where M1 cells stratify. Thus, to ensure effective data collection, we optically recorded the glutamate release onto labeled dendrites of selected cells, and post hoc reconstructed and identified the cells based on 2-photon z-stacks. The glutamate released onto OFF-IPL dendrites labeled in the Opn4^Cre/+^ mouse was sustained and intensity-encoding (Fig. 6b). Many of these cells exhibited too dense dendritic arbors to be considered M1 cells. Instead, these cells may correspond to M3 ipRGCs or to the bistratified RGCs, which as our SBEM analysis demonstrates, receive ectopic synapses (Extended Data Fig. 3). Nonetheless, the observed light-evoked glutamate release at the far OFF-IPL (where the axon terminals of OFF-BCs stratify) demonstrates that ectopic synapses transmit sustained intensity-encoding light information.

**Figure 6.**
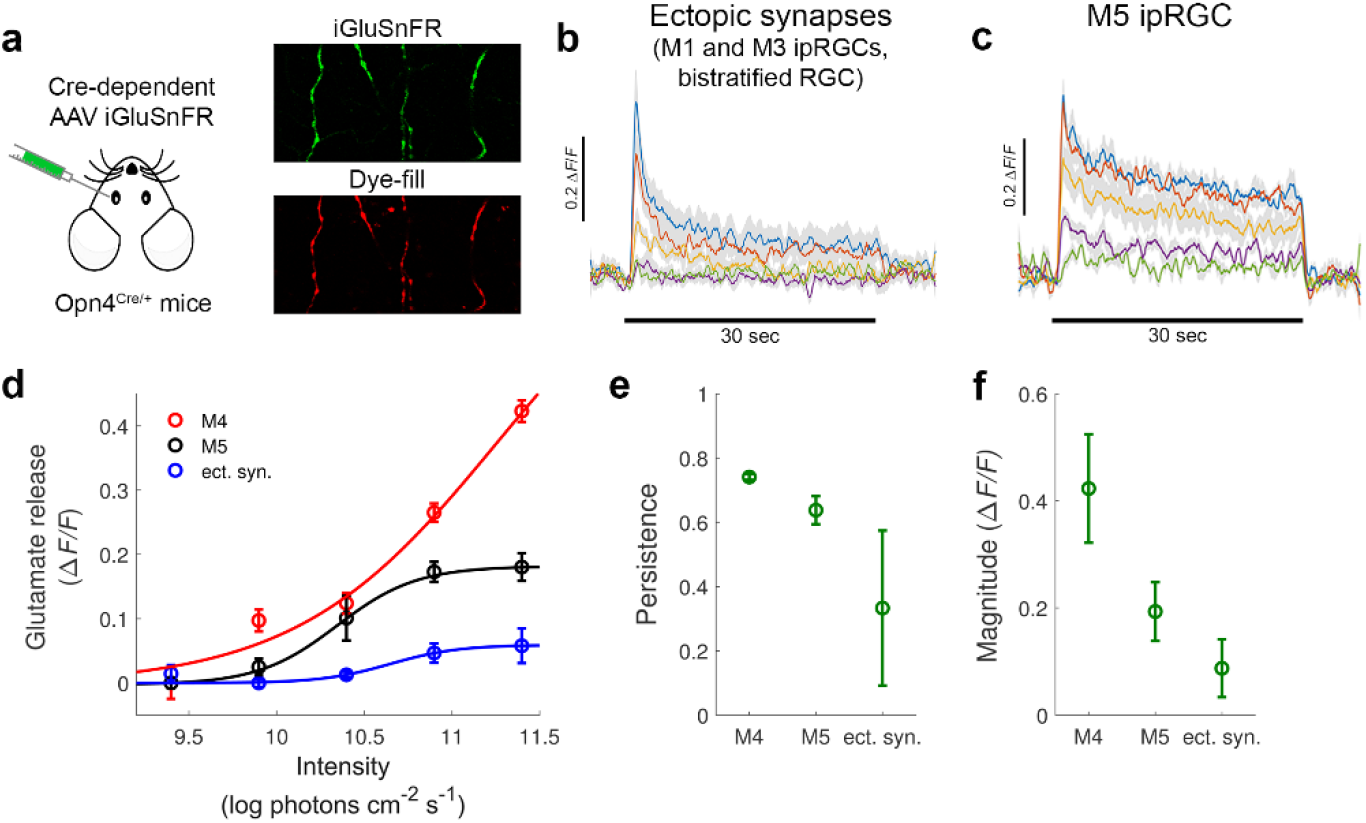
Sustained intensity-encoding glutamate release at ectopic synapses. **a**, Glutamate release at ectopic synapses and onto ipRGCs was measured following intravitreal injections of Cre-dependent iGluSnFR AAVs into the eyes of Opn4^Cre/+^ mice that selectively label melanopsin-expressing ipRGCs. Ectopic synapses were targeted by imaging iGluSnFR-positive dendrites in the OFF IPL. To target M4 and M5 cells, selected iGluSnFR-positive cells were patched with red dye in the pipette, and their arbor, identified by the red dye fill, was imaged. **b**,**c**, Glutamate release (Δ*F*/*F*) at ectopic synapses (n = 9) and onto M5 ipRGCs (n = 5). Glutamate release was measured in response to 30 sec of light (indicated by a horizontal black bar) and at 5 different light levels (log photons cm^-2^ s^-1^; depicted in different colors). **d**, Steady-state glutamate release as a function of stimulus intensity for M4 (n = 5, raw data presented in Fig. 4) and M5 (n = 5) ipRGCs, and ectopic synapses (n = 9), ect. syn.). Steady-state response was estimated as the mean response over the 5 last seconds of the light phase of the stimulus, and fitted to the Naka-Rushton function (symbols, mean; error bars, s.e.m). **e**, Persistence of glutamate release for M4, M5, and ectopic synapses. Persistence (mean±s.d.) of glutamate release through ectopic synapses is slightly lower than that onto M4 and M5. **f**, Steady-state magnitude of glutamate release in response to the highest light intensity was greatest for M4 cells, lower for M5 cells, and lowest for ectopic synapses. (mean±s.d.).

To compare the kinetics of glutamate release between ectopic monad synapses and terminal dyad synapses, we compared glutamate release at ectopic synapses to the glutamate release onto M4 and M5 ipRGCs that receive their BC input only through terminal dyad synapses in the ON-IPL (65%-75% IPL depth). Glutamate released onto M5 cells was measured similarly to that onto M4 cells. Figure 6d-f shows how the capacity for intensity encoding varied among M4 and M5 ipRGCs and ectopic synapses. These results demonstrate that ectopic monad synapses are capable of transmitting sustained intensity-encoding signals, but at a somewhat reduced capacity than terminal dyad synapses.

### Selectivity between bipolar cells and retinal ganglion cells

Our SBEM data allowed us the shed light on the developmental processes that underlie the selectivity between RGCs and BCs. We asked whether we could predict the bipolar input based only on the stratification profiles of RGCs and BCs, and the terminal field area of BCs (Figure 7a). Unexpectedly, this very simple model could predict the BC-RGC selectivity for some RGCs but not for others. The selectivity between BCs and examined ipRGCs could be predicted solely by their stratification depth and the terminal field area of BCs. In other words, BCs provided input to ipRGCs that costratified with them, with no obvious selectivity. This was true for all ipRGC except for M1 ipRGCs. Despite stratifying in the far OFF-IPL, M1 cells received input only from ON-BCs, predominantly from Type 6 BCs, and avoided axon terminals from OFF-BCs (Figure 7b-d, left column). As for ipRGCs, the selectivity between BCs and examined conventional RGCs could also be predicted using our simple model. (Figure 7b-d). This was true for the ON-OFF-DSGC, OFF *α*RGC, and the newly identified bistratified RGC that receives ectopic synapses (compare Figure 7b and c). However, closer inspection of BC input patterns to RGCs revealed fine variations that could not be explained by the cells’ stratification depth, especially in the case of ON-OFF-DSGC (Figure 7d). These results suggest that the connectivity between BCs and certain ipRGC and conventional RGC types rely on factors other than stratification depth.

**Figure 7.**
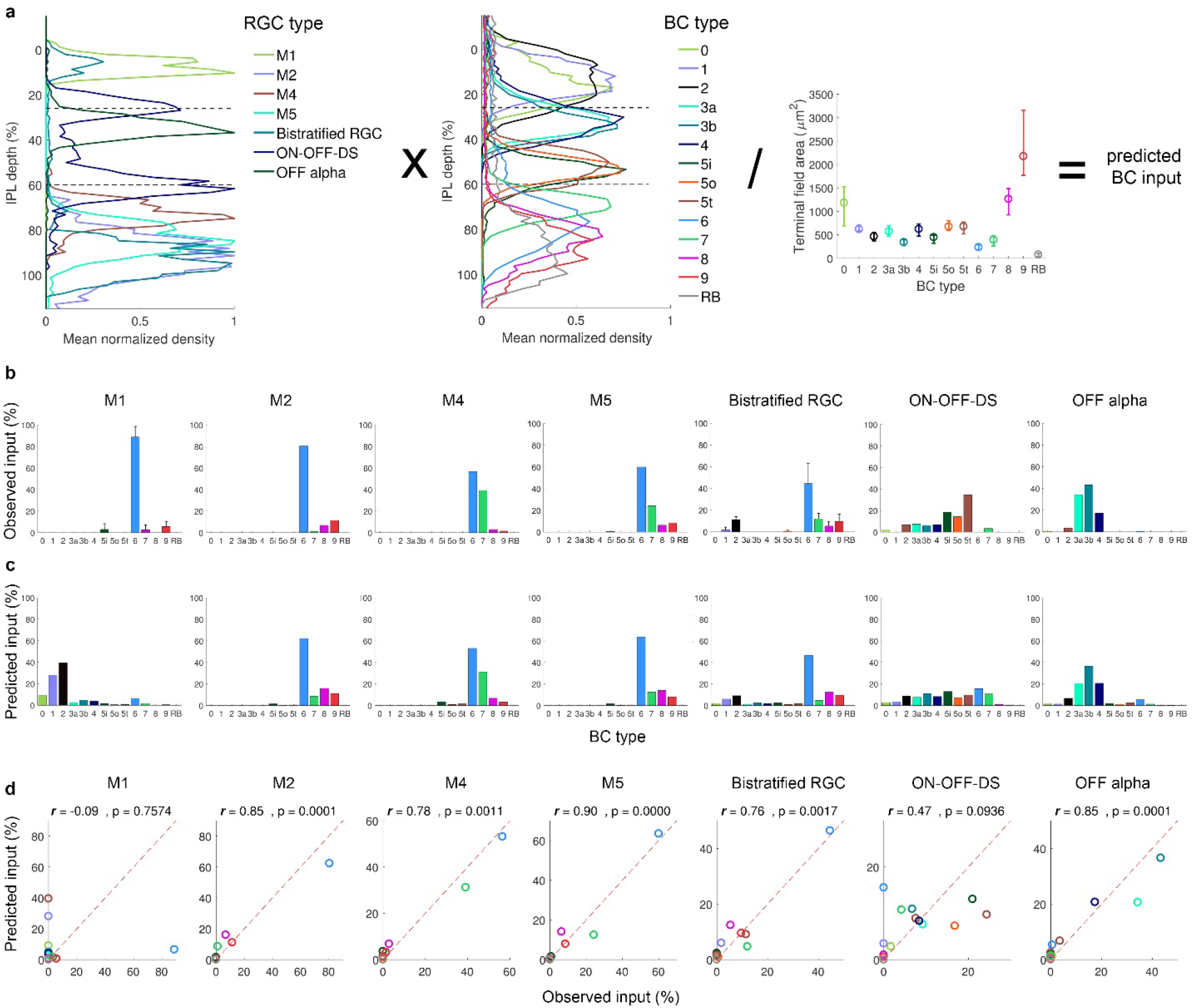
Stratification depth determines the selectivity between BCs and some RGCs. **a**, Input patterns from BCs to RGCs can be predicted by their stratification depth and the terminal field area of BCs. **b**, Observed BC input to M1-M5 ipRGCs, a bistratified RGC, ON-OFF-DSGC, and an OFF alpha RGC. **c**, Predicted input of BCs to RGCs. **d**, Predicted vs. observed BC input to RGCs. Red dashed line depicts the identity (predicted = observed) line. The Spearman correlation coefficient and associated p-value are indicated for the different RGC types.

## Discussion

Among retinal output neurons, only ipRGCs stably encode environmental light intensity, enabling diverse light-dependent physiological effects. We have shed new light on the role of bipolar inputs in this unique capacity of ipRGCs. We describe which bipolar types synapse on ipRGCs, what their synapses look like, and how the capacity of their output signals to encode intensity differ from those driving other RGC types. We conclude that ipRGCs do sample from specialized BCs and, in some cases, at specialized synapses, but that post-synaptic mechanisms also contribute to the special capacity for intensity encoding.

We showed that ipRGCs receive their synaptic input only from ON-BCs, predominantly from BC Types 6 and 7, through either ectopic monad synapses (in the OFF-IPL) or terminal dyad synapses (in the ON-IPL). In general, ipRGCs receive synaptic input from BCs that costratify with them, with no obvious selectivity. However, OFF-stratifying M1 ipRGCs receive their synaptic input almost exclusively through ectopic synapses and largely avoid axon terminals from OFF-BCs. Such selectivity is crucial for the ability of M1 cells to convey ON light information, which is the main drive for the pupillary light reflex and the photoentrainment of our circadian clock ^33^. Ectopic synapses are in fact ubiquitous in the OFF-IPL, and appear to form a dedicated circuit with ipRGCs and their network amacrine cells.

Bipolar inputs to ipRGCs derive mainly from Types 6 and 7, through either ectopic monad synapses or conventional dyad synapses. Glutamate released at these two types of synapses is sustained and intensity encoding. Yet, the two types of synapse are markedly different. Dyads have only one ribbon while monads contain several and up to a dozen or more ribbons. Considering that ribbon synapses facilitate sustained neurotransmitter release at the synaptic cleft^34,35^, it would be important to compare systematically the kinetics of both synapse types.

Functional imaging confirmed sustained, intensity-encoding glutamate release onto ipRGC dendrites. This indicates that bipolar Types 6 and/or 7 are capable of such signaling but, surprisingly, this was not unique to them. The glutamate signal was also sustained and intensity-encoding onto each of a variety of conventional RGCs, and throughout the depth of the IPL, including the OFF sublayer. The stark differences in intensity encoding between ipRGCs and conventional RGCs reflects to a large degree differences in postsynaptic temporal filtering of intensity signals. Nonetheless, bipolar types did differ in the strength of their intensity signal, intensity-encoding did vary modestly with IPL depth, suggesting characteristic differences among. Additionally, the effect of GABAergic and glycinergic feedback inhibition on the persistence of BC glutamate release was minimal, if any, consistent with previous a report^36^. Together, these results demonstrate that the key locus for temporal filtering of sustained intensity-encoding signals is postsynaptic to the bipolar axon terminal, perhaps through modulations of glutamate receptor kinetics, intrinsic membrane properties or feed-forward inhibition. The variation in persistence and steady-state magnitude among bipolar output signals could have multiple sources, but almost certainly include cell-type-specific variation in metabotropic glutamatergic signaling in ON bipolar cells, axon terminal biophysics and vesicular glutamate release ^36-38^.

The observed intensity-encoding glutamate release across the IPL, even in the OFF-IPL, is surprising. Indeed, vGluT3 ACs, whose dendrites span the middle of the IPL between the ON and OFF ChAT bands, also use glutamate as their neurotransmitter ^39^. Thus, the measurements of glutamate released at IPL depths 35%-55% likely reflects also the contribution of vGluT3 transmission, in addition to BC transmission. Moreover, measurements of the glutamate release within the IPL represent the average glutamate released onto the dendrites of many RGCs and ACs, and it is not indicative of the input onto single RGCs. Our measurements of the glutamate release onto the dendrites of identified ipRGCs and conventional RGCs addressed this limitation.

While ipRGCs transmit intensity-encoding signals, conventional RGCs are optimized for the transmission of contrast-encoding signals and support detection and recognition of objects, color and motion^3^. Though contrast and intensity signals are carried through these distinct sets of retinal outputs, they also appear to interact in the context of light adaptation, which allows the retina to encode contrast over a large range of light intensities. ipRGCs are known to inject intensity signals back into retina by means of chemical synapses onto dopaminergic ACs^40^. This intraretinal dopaminergic signaling, in turn, modulates chemical and electrical synapses and modifies the functional properties of retinal neurons, thereby modulating the dynamic range over which the retina operates^41^. Additionally, ipRGCs have been recently shown to inject their intensity signals by means of electrical transmission through gap junctions to a family of spiking polyaxonal amacrine cells whose somata are unconventionally placed in the ganglion cell layer^30,42^. However, it is unclear whether these intensity signals are transmitted within the retina or modulate its sensitivity. Supplementary Information is available in the online version of the paper.

## Acknowledgements

We thank Alex kolodkin for kindly providing us with the Pcdh9^Cre/+^ mice. Many colleagues provided invaluable theoretical, data-analytic and technical advice. They included Jim McIlwain, Bart Borghuis, Geoff Williams, Scott Cruikshank, and Shane Crandall. Dianne Boghossian maintained the mouse colony and genotyped experimental mice. We thank Maureen Estevez Stabio for kindly providing us with a confocal z-stack for an M5 ipRGC. We thank Loren L. Looger, Janelia Research Campus, Howard Hughes Medical Institute for sharing their iGluSnFR glutamate indicator. This project was supported by the Banting Postdoctoral Fellowship of Canada to S.S., The Sidney A. Fox and Dorothea Doctors Fox Postdoctoral Fellowship in Ophthalmology and Visual Sciences to S.S., and NIH grant (R01 EY12793) and an Alcon Research Institute Award to D.M.B.

## Author Contributions

S.S. and D.M.B. designed the study and developed the theoretical framework. S.S. performed all imaging, electrophysiological recordings, and intracellular dye-fills. S.S., E.K., G.M., M.B. and T.Z. performed intravitreal injections. S.S., D. M. B., C.P., E.K., M.B., C.E., A.N.G., L.C., M.T.K., G.M., A.B.L., T.Z., D.S. and H.D. traced and classified retinal neurons from a serial block-face electron microscopy data set and light microscopic depth series, and performed immunostaining of retinas. S.S. analyzed all morphological, electrophysiological, and functional imaging data. S.S. and D.M.B. wrote the paper.

Correspondence and requests for materials should be addressed to

## Methods

### Animals

All procedures were in accordance with National Institutes of Health guidelines and approved by the Institutional Animal Care and Use Committee at Brown University. We used adult (2 - 2.5 months old; either sex) wildtype C57BL/6J mice (Jackson Laboratory) and the melanopsin reporter strain Opn4cre/+ 20 that marks M1-M5 ipRGCs, donated by Dr. Samer Hattar, Johns Hopkins University.

### Intravitreal injections of glutamate florescent indicator

Mice (C57BL/6J or Opn4^cre/+^) were anesthetized with isoflurane (3% in oxygen; Matrx VIP 3000, Midmark). Two viral vectors inducing expression of the glutamate indicator iGluSnFR were used (Vector Core, UPenn; 1.5 –2 μl of ~10^12^ units/ml): a Cre-dependent vector (AAV1.CAG.Flex.iGluSnFR) and a non-specific one (AAV1.hSyn.iGluSnFR). Viral vectors were injected into the vitreous humor of the right eye through a glass pipette using a microinjector (Picospritzer III, Science Products GmbH). Animals were killed and retinas harvested 14-21d later. iGluSnFR was expressed mainly in RGCs and amacrine cells of the ganglion cell layer.

### Tissue harvest and retinal dissection

Eyes were removed and immersed in oxygenated Ames medium (95% O_2_, 5% CO_2_; Sigma-Aldrich; supplemented with 23 mM NaHCO_3_ and 10 mM D-glucose). Under dim red light, the globe was cut along the ora serrata, and cornea, lens and vitreous removed. Four radial relieving cuts were made in the eyecup. The retina was flat-mounted on a custom-machined hydrophilic polytetrafluoroethylene membrane (cell culture inserts, Millicell; ^43^) using gentle suction, and secured in a chamber on the microscope stage. Retinas were continuously superfused with oxygenated Ames’ medium (32–34°C).

### Two-photon functional glutamate imaging

Glutamate imaging and whole-cell recordings were conducted on a multiphoton Olympus FV1200MPE BASIC (BX-61WI) microscope equipped with a 25x, 1.05 NA water-immersion objective (XLPL25XWMP, Olympus) and an ultrafast pulsed laser (Mai Tai DeepSee HP, Spectra-Physics) tuned to 910 nm. Epifluorescence emission was separated into “green” and “red” channels with a 570 nm dichroic mirror and a 525/50 bandpass filter (FF03-525/50-32, Semrock, green channel) and 575-630 nm bandpass filter (BA575-630, Olympus, red channel), respectively. The microscope system was controlled by FluoView software (FV10-ASW v.4.1). Images of 256 × 128 pixels representing to 84 × 42 μm on the retina were acquired at 15 Hz (zoom setting of 6).

We pharmacologically blocked various receptors and pathways with one or more chemicals: 1) L-AP4 (a group III metabotropic glutamate receptor agonist; acting on the metabotropic glutamate receptor mGluR6); 2) picrotoxin (a non-competitive channel blocker for GABA_A_ receptors); or 3) strychnine (an antagonist of glycine receptors). All were purchased from Tocris.

### Visual stimulation

Patterned visual stimuli, synthesized by custom software using Psychophysics Toolbox under Matlab (The MathWorks), were projected (AX325AA, HP) and focused onto photoreceptor outer segments through the microscope’s condenser ^22^. The projected display covered 1.5 × 1.5 mm; each pixel was 5×5 μm. The video projector was modified to use a single UV LED lamp (NC4U134A, Nichia). The LED’s peak wavelength (385 nm) shifted to 395 nm after transmission through a 440 nm short-pass dichroic filter (FF01-440/SP, Semrock), a dichroic mirror (T425lpxr, Chroma), and various reflective neutral density filters (Edmund Optics).

Quantum catches were derived from the stimulus spectrum (measured using an absolute-irradiance-calibrated spectrometer [USB4000-UV-VIS-ES, Ocean Optics]) and spectral absorbances of mouse rod, cone, and melanopsin pigments ^4,44^. Quantum catches were very similar among rods, cones, and melanopsin (~11.9 log photons cm^-2^ s^-1^ at the highest light stimulus intensity), independent of the cones’ relative expression of S- and M-cone pigments ^45,46^.

To study the cells’ ability to encode increments in the absolute irradiance level, we used bright circular spots on a dark background (diameter=200 μm, Michelson contrast=0.95, stimulus duration=30 sec, inter-stimulus duration=15 sec, 6 repetitions) at 5 irradiance levels at the plane of the photoreceptors (9.4-11.4 log photons cm^-2^ s^-1^). To allow adaptation of the photoreceptors to the scanning laser, the data from the first repetition (out of 6) in each trial was excluded from analysis. To study the ability of individual cells to encode decrements in the absolute irradiance level, we used dark circular spots of five intensities (6-10.4 log photons cm^-2^ s^-1^) on a bright background (12.5 log photons cm^-2^ s^-1^). Whereas, to study the glutamate release across the entire in response to light decrements, we used dark circular spots of five intensities (5.5-9.9 log photons cm^-2^ s^-1^) on a bright background (11.9 log photons cm^-2^ s^-1^). To assess directional tuning, we used a sinusoidal grating spanning two spatial periods (spatial frequency=0.132 cycle/degree, Michelson contrast=0.95, stimulus duration=3.65 sec, inter-stimulus duration=5 sec at uniform mean grating luminance) drifted in 8 randomized directions (45° interval, drift speed=4.5 degree/sec, 4 repetitions). Frames of the stimulus movie appeared for 50 μs during the short 185 μs interval between successive sweeps of the imaging laser; thus, no stimulus was presented during the interval of laser scanning and associated imaging (300 μs / sweep). The very rapid stimulus flicker (>2000 Hz) was well above critical fusion frequency in mice ^47^.

### Patch recording and dye filling of ganglion cells

Whole-cell patch-clamp current-clamp and voltage-clamp recordings of isolated flat-mount retinas were performed as described^48^, using a Multiclamp 700B amplifier, Digidata 1550 digitizer, and pClamp 10.5 data acquisition software (Molecular Devices; 10 kHz sampling). Pipettes were pulled from thick-walled borosilicate tubing (P-97; Sutter Instruments); tip resistances were 5–6 MΩ when filled with internal solution, which, for current-clamp recordings, contained (in mM): 120 K-gluconate, 5 NaCl, 4 KCl, 2 EGTA, 10 HEPES, 4 ATP-Mg, 7 phosphocreatine-Tris, and 0.3 GTP-Tris, pH 7.3, 270–280 mOsm). For voltage-clamp recordings, the internal solution contained (in mM): 120 Cs-methanesulfonate, 5 NaCl, 4 CsCl, 2 EGTA, 10 HEPES, 4 ATP-mg, 7 Phosphocreatine-tris, and 0.3 GTP-tris, pH 7.3, 270–280 mOsm). Red fluorescent dye (Alexa Fluor 568; Invitrogen) was added to the pipette for visual guidance under two-photon imaging.

### Analysis of glutamate imaging and electrophysiological data

Acquired time-series images were registered, and dendritic glutamate responses were analyzed using FluoAnalyzer ^22^ and custom Matlab scripts^48^. We employed two distinct approaches. To estimate the glutamate release at different depths across the IPL, the whole imaged field (84 × 42 μm) was taken as the region of interest (ROI). Whereas, to estimate the glutamate release onto dendrites of a particular RGC, we patched-filled the RGC with a red fluorescent dye, and imaged fields that included dendrites showing both red (filled cells) and green (iGluSnFR) fluorescence. Later, to identify possible synapses on the targeted RGC’s dendrites, we scanned the dendrites for areas where the standard deviation fluorescence over a time series of iGluSnFR responses was high. Such areas appeared as hot spots on the dendrites which were then manually selected as ROIs (7-10 ROIs per imaged field).

The space-averaged pixel intensity within such ROIs was the activity readout for the associated cell, a proxy for its spike rate^49^. Fluorescence responses are reported as normalized increases as follows:

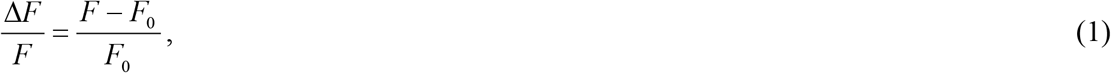

where *F* denotes the instantaneous fluorescence and *F*_0_ the mean fluorescence over a 1-second period immediately preceding stimulus onset.

Irradiance-Response (IR) curves were calculated using the steady-state response, which was taken as the 5 last sec of the 30 sec light-evoked response. The ability of cells to report the absolute irradiance was assessed by fitting the sigmoidal Naka-Rashton function ^50^ to the cell’s steady state response to various stimulus irradiance levels (*R*):

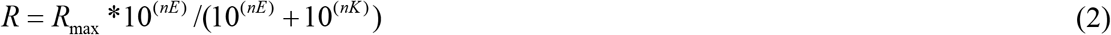

Where *R*_max_ stands for the cell’s predicted maximum response, *n* stands for the slope of the function, *E* stands for the irradiance measured in units of log photons cm^-2^ s^-1^, and *K* represents for the cell’s sensitivity.

The magnitude of steady state glutamate release was calculated as the difference between the maximum and minimum glutamate release encountered across the five stimulus irradiance levels tested. This magnitude is influenced by the permanence of the response and by the capacity of glutamate release to encode irradiance. Magnitude of zero indicates no variation in the steady state response across stimulus irradiance, whether because glutamate release is transient and/or because glutamate release is sustained but not correlated with intensity.

To quantify how sustained the glutamate release is in relative terms, we developed the persistence index:

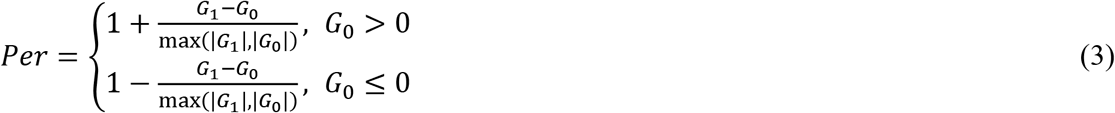

where *G*_0_ represents the average glutamate release over the first 5 sec of the stimulus, and *G*_1_ represents the average glutamate release over the last 5 sec of the stimulus of the highest-irradiance stimulus. A persistence index of 0 indicates that the glutamate signal returned the baseline by the termination of the stimulus (a perfectly transient response). Whereas, a persistence index of 1 indicates that the glutamate signal remained constant throughout the stimulus duration (a perfectly sustained response).

To calculate the sensitivity of glutamate release we fit a third-order polynomial to the IR curve, and the threshold irradiance that corresponded to a fixed response criterion was interpolated; sensitivity was estimated as the reciprocal of this threshold irradiance ^51^. The response criterion was set to a value low enough (0.022 Δ*F*/*F*) such that it was represented in IR curves across the whole IPL. For excitatory currents recordings, we used a response criterion of 10 pA.

The dynamic range of steady state responses was calculated as the 10^th^ and 90^th^ percentiles of the first derivative of the fit Naka-Rashton function. The fit of the Naka-Rashton function did not reach saturation at 85% IPL depth; this dynamic range estimate was deemed unreliable and therefore was excluded.

For all stimuli tested, the response amplitude represented the normalized fluorescence increases (for glutamate imaging), average firing rate (for current-clamp recording under control conditions), average lower envelope of the voltage response (for current-clamp recording under any pharmacological manipulation), or average current response (for voltage-clamp recording under any condition). All data were analyzed using custom Matlab scripts.

To evaluate the glutamate release onto individual RGC dendrites at the ON-OFF transition zone, we selected ~150 regions of interest (ROIs) from the fluorescence time series acquired in response to the stimuli of the highest light intensity. ROIs were classified as either dominated by an ON or OFF response based on whether the mean response during the stimulus’ light phase was higher (‘ON ROIs’) or lower (‘OFF ROIs’) than that during the stimulus’ dark phase. Thereafter, the mean response of ON and OFF ROIs was calculated. This procedure was repeated for several IPL depth profiles.

The preferred direction of a cell was estimated as the angle of the vector sum following ^35^:

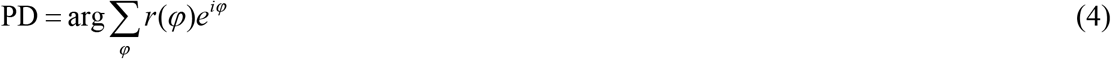

where *r* is the response amplitude to stimuli moving at direction *φ* (0, 45,…,315). The direction selectivity index (DSI) of cells which may range between 0 (no direction selectivity) and 1 (highest direction selectivity) was calculated as:

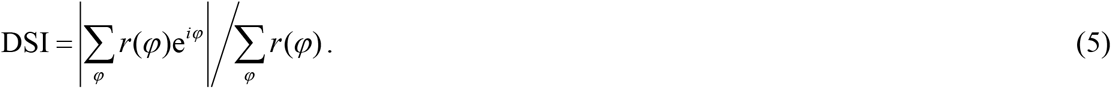

The response amplitude *r* represented the average response from the stimulus onset to 2 sec following the termination of the stimulus, to capture the OFF responses.

### Immunohistochemistry of retina and brain

After recording, retinas were fixed (4% paraformaldehyde, 30 min, 20°C) and counterstained with one or more antibodies: 1) guinea pig anti-Rbpms (1:1000, RNA-binding protein with multiple splicing; 1832-Rbpms, PhosphoSolutions), a pan-ganglion-cell marker ^52^; 2) goat anti-ChAT (1:100, anti-choline acetyltransferase; AB144P, Millipore); 3) rabbit anti-melanopsin (1:1000, Advanced Targeting Systems); or 4) chicken anti-GFP (1:1000, ab13970 Abcam), to enhance the fluorescence of the GFP-based GCaMP6f indicator.

### Reconstruction of retinal neurons based on a serial electron microscopy data set

Tissue preparation and EM acquisition were performed as previously described ^28^. In short, retina (k0725) from an adult wild-type mouse (C57BL/6; postnatal day 30) was stained for EM while preserving the intracellular structure and details. A retinal block face of ~200 × 400 μm was imaged using a serial block-face scanning electron microscope system. The incident electron beam had an energy of 2.0 keV and a current of ~110 pA. Images were acquired with a pixel dwell time of 2.5 μs and size of 13.2 nm × 13.2 nm. The section thickness was set to 26 nm. 10,112 consecutive block faces were imaged, resulting in aligned data volumes of 4,992 × 16,000 × 10,112 voxels, corresponding to an approximate spatial volume of 50 × 210 × 260 μm^3^. The retinal imaged region spanned the IPL and included parts of the GCL and INL. To facilitate viewing in KNOSSOS (http://www.knossostool.org), the data set was split into cubes (128 × 128 × 128 voxels).

The skeletons of cells were manually traced using the KNOSSOS annotation platform. Skeletons were annotated by placing nodes in the relative center of a neurite’s section, branch points, and somas. All skeletons were traced by at least two observers and any discrepancies resolved to ensure accuracy. We assigned bipolar cells to established categories ^11,29,53^ based on stratification level and lateral dimensions of the axonal arbor. Where we had dense sampling of neighboring bipolar cells, the tiling pattern of arbors further aided the type assignment.

For each BC, we calculated the density of neurites as a function of IPL depth, where IPL depth=0% represent the INL-IPL boundary, and IPL depth=100% represent the IPL-GCL boundary. The 5^th^ percentile neurite density of the axon terminal of all type 2 BCs marked the INL-IPL boundary, whereas the 95^th^ percentile neurite density of the axon terminal of all rod BCs marked the IPL-GCL boundary. The slight tilt of the retinal laminae relative to the cutting plane was accounted for when calculating the IPL depth of cells. All analyses on skeleton data were performed using MATLAB.

Morphological statistics of EM traced cells were compared to those of traces obtained from light microscopy (LM) data. EM traces of the dendritic/axonal arbors of several RGCs and ACs were incomplete due to the small size of the EM volume. Thus, we gave priority to morphological parameters that are least susceptible to the incompleteness of traces. These were the stratification pattern, soma diameter, branch point density, and number of primary dendrites.

### Statistical analysis

To test statistically the effect of transmission stage (glutamate release, excitatory currents, and firing rate) on the persistence and magnitude of responses we used a permutation ANOVA. To test the persistence and magnitude of responses between M4 ipRGCs and conventional ON-DSGCs, we used a permutation t-test. These data often violated both the normality (Kolmogorov Smirnov test) and homoscedasticity assumptions (Bartlett’s test). Therefore, we utilized appropriate permutation tests with a significance level of 0.05. All analyses performed in the R statistical software.

---

**Extended Data Figure 1.**
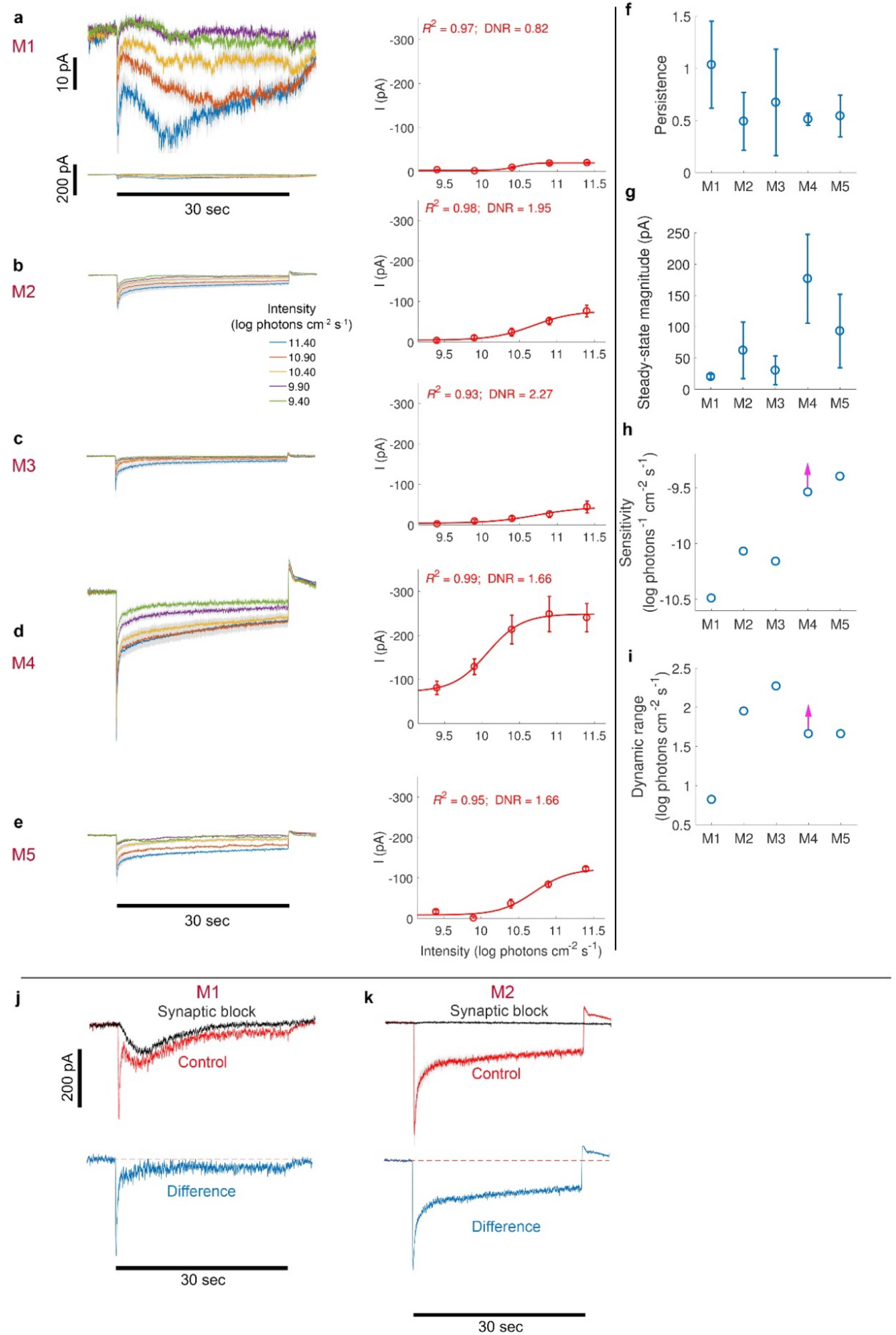
ipRGCs differ in properties of synaptic excitatory input, but all receive sustained and intensity-encoding input. **a (left, top)**, Excitatory currents in M1 ipRGCs (*n* = 4) in response to five stimulus light intensities (depicted in different colors). Horizontal black bar below the traces indicates the timing of the 30-s stimulus (spot diameter, 200 μm). Inward currents in M1 cells comprise two phases: an initial transient phase dominated by rods/cones synaptic input, and a later sustained phase dominated by the intrinsic photosensitivity of melanopsin. The higher the intensity, the larger the amplitude of the second phase is. Blocking synaptic transmission using L-AP4, D-AP-5, DNQX, and ACET eliminated the initial transient phase (**j**). The difference curve reveals a small sustained component of synaptic input. **a (left, bottom)**, Similar excitatory currents plotted at a scale similar to the reminder of the ipRGCs types (**b**-**e**). **a (right)**, Intensity-response (IR) curve based on data presented in **a (left).** Data points and error bars indicate the average±SEM of the steady state response, assessed during the last 5 s of the stimulus. Red line denotes the fitted sigmoidal Naka-Rushton function. The coefficient of determination (*R*^2^) for the fit is indicated. Light-evoked excitatory currents in M1 are sustained and intensity-encoding, but of low amplitude. **b-e**, Similar plots for M2 (*n* = 6) (**b**), M3 (*n* = 5) (**c**), M4 (*n* = 5) (**d**), and M5 (*n* = 4) (**e**) ipRGCs. Note that excitatory currents in M2-M5 ipRGCs returned to baseline, or in some instances, even over shoot abruptly at light off. In contrast, excitatory currents in M1 ipRGCs leveled off gradually at light off, indicating a dominant contribution of the melanopsin photoresponse only in M1 ipRGCs. Indeed, blocking synaptic transmission in an M2 cell completely eliminated the response (**k**). **f**, The persistence of excitatory currents in response to the highest light intensity, calculated as the ratio between the response over the last and first 5 sec of stimulus (persistence of 0 or 1 indicates a complete transient or sustained response). Persistence typically ranged 0.5 – 1 for M2-M5 ipRGCs. M1 ipRGCs showed persistence larger than 1, indicating that the response during the initial synaptically-driven phase of response was smaller than that during the subsequent melanopsin-driven phase. **g**, The steady-state magnitude of excitatory currents in response to the highest light intensity differed considerably across ipRGC types, with M4 ipRGCs showing the largest magnitude. **h**, Sensitivity of the current responses in all ipRGC types. Sensitivity was calculated by fitting a third-order polynomial to the IR curve, and the threshold intensity that corresponded to a fixed response criterion was interpolated (see Methods); sensitivity was estimated as the reciprocal of this threshold intensity. Sensitivity generally decreased when moving from M1 to M5 ipRGCs. **i**, Dynamic range of sensitivity based on the current responses in all ipRGC types. The dynamic range of steady state responses was calculated as the 10^th^ and 90^th^ percentiles of the first derivative of the fit Naka- Rashton function. M3 and M1 ipRGCs showed the largest and smallest dynamic ranges, respectively.

**Extended Data Figure 2.**
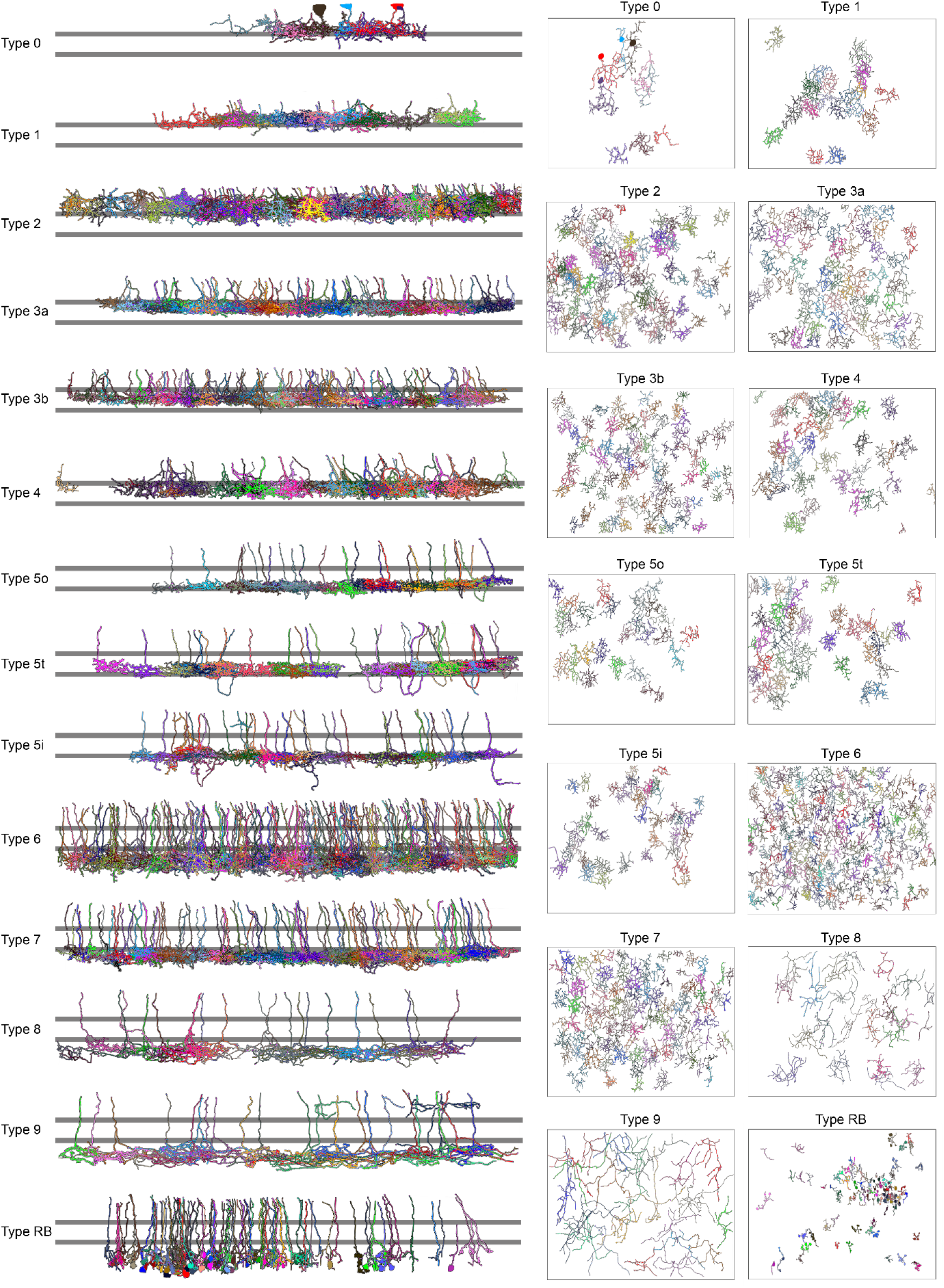
All bipolar cells, separated by type, reconstructed in this study. The different BC types in the plane of the retina (right), and at an orthogonal view (left), relative to the ON and OFF ChAT bands (two horizontal gray stripes).

**Extended Data Figure 3.**
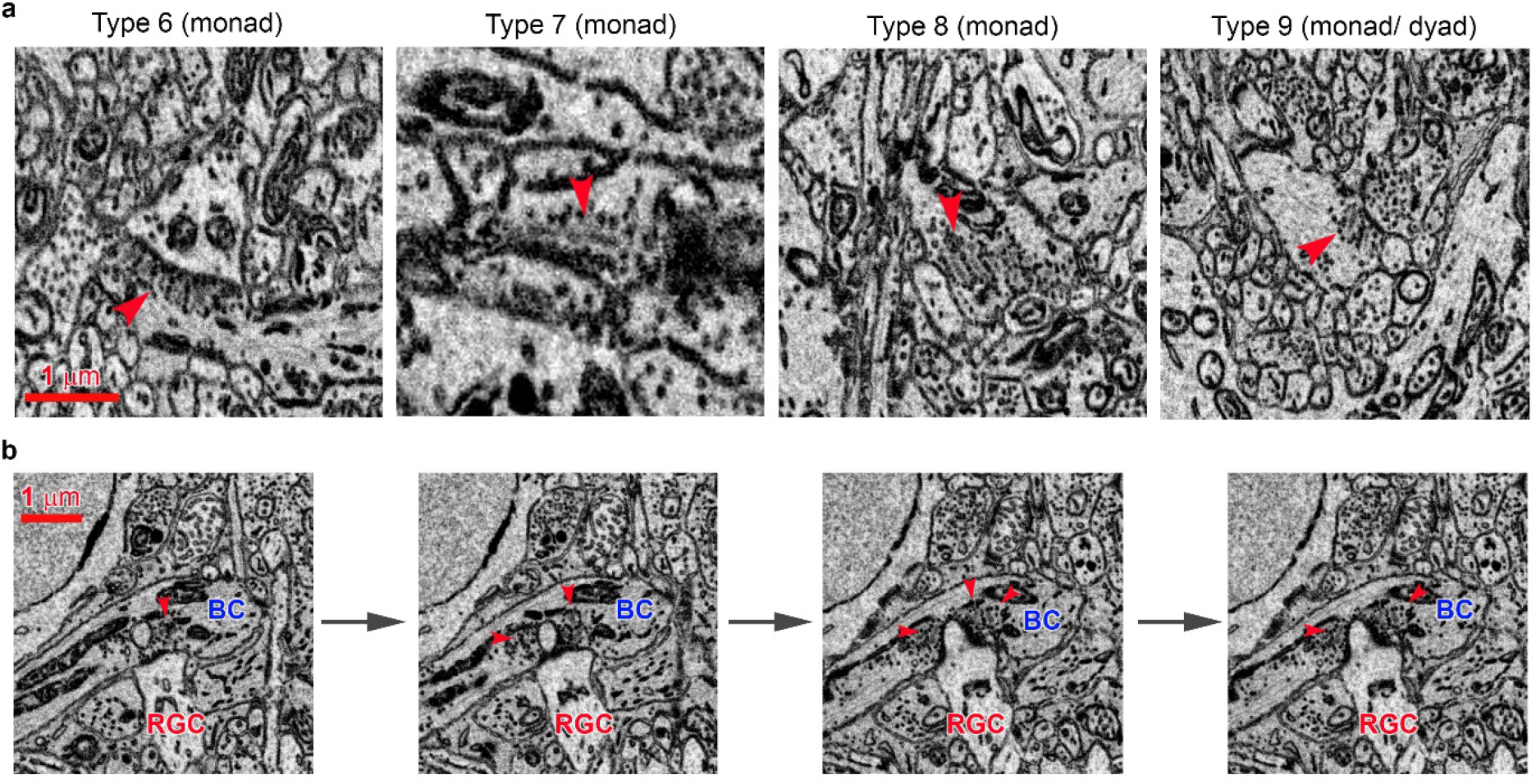
Diversity of ectopic synapses. **a**, Example ectopic synapses encountered on the axonal shafts of BC Types 6, 7, 8, and 9. Red arrow points to synaptic ribbons and aggregate of neurotransmitter vesicles located at the membrane of presynaptic BCs. **b**, A time series of electron micrographs of a monad synapse (membrane swolling) exhibiting rings of neurotransmitter vesicles around several ribbons.

**Extended Data Figure 4.**
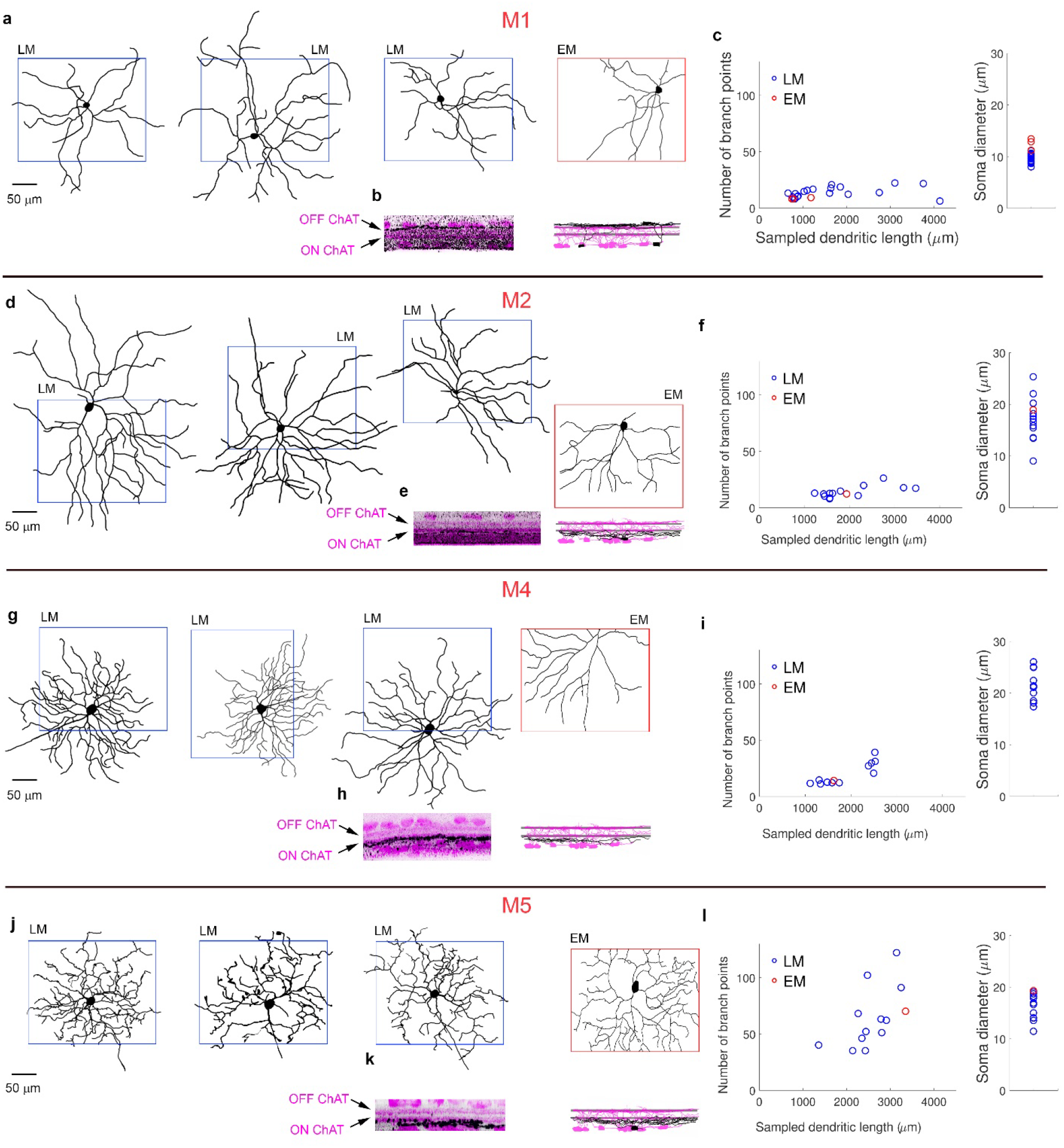
Morphological comparison between SBEM-reconstructed and genetically identified ipRGCs. **a.** Comparison of selected morphological statistics between SBEM-reconstructed (EM) presumptive M1 ipRGCs and M1 cells identified using light microscopy (LM). Three left traces (enclosed by a blue rectangle) represent the *en face* projections of three M1 cells based on depth series (z-stacks) of confocal light micrographs. Fourth trace from the left (enclosed by a red rectangle) represents a presumptive M1 cell identified in the SBEM volume. **b**, **left**, Orthogonal projection of a representative LM-reconstructed cell (black) overlaid with anti-ChAT immunostaining (magenta). **b**, **right**, Orthogonal projection of a representative EM-reconstructed cell (black) overlaid with reconstructed ON-OFF DSGCs whose dendritic arbors roughly mark the two ChAT bands formed by the processes of ON and OFF starburst amacrine cells (magenta). The stratification profile of the LM- and EM-reconstructed cells are similar. **c**, **left**. Comparison of the number of branch points as a function of sampled dendritic length for LM-reconstructed (blue) and EM-reconstructed (red) cells. **c**, **right**. Comparison of the soma size between LM-reconstructed (blue) and EM-reconstructed (red) cells. Note that the distal dendrites of EM reconstructed cells were often clipped due to area limitations of the SBEM dataset, which may render comparison between EM and LM traces misleading. To overcome this, we clipped LM traces using a rectangular frame the size of the SBEM volume (260 μm × 210 μm). Thus, both EM and LM traces sampled an equivalent area of a cell’s dendrosomatic profile. To avoid bias in the location of the sampling frame, we repeated the sampling process 2-3 times, each time sampling a different part of the dendritic arbor(s). **d**-**f**, The same as **a-c** but for M2 ipRGCs. **g**-**i**, The same as **a-c** but for M4 ipRGCs. **j**-**l**, The same as **a-c** but for M5 ipRGCs.

**Extended Data Figure 5.**
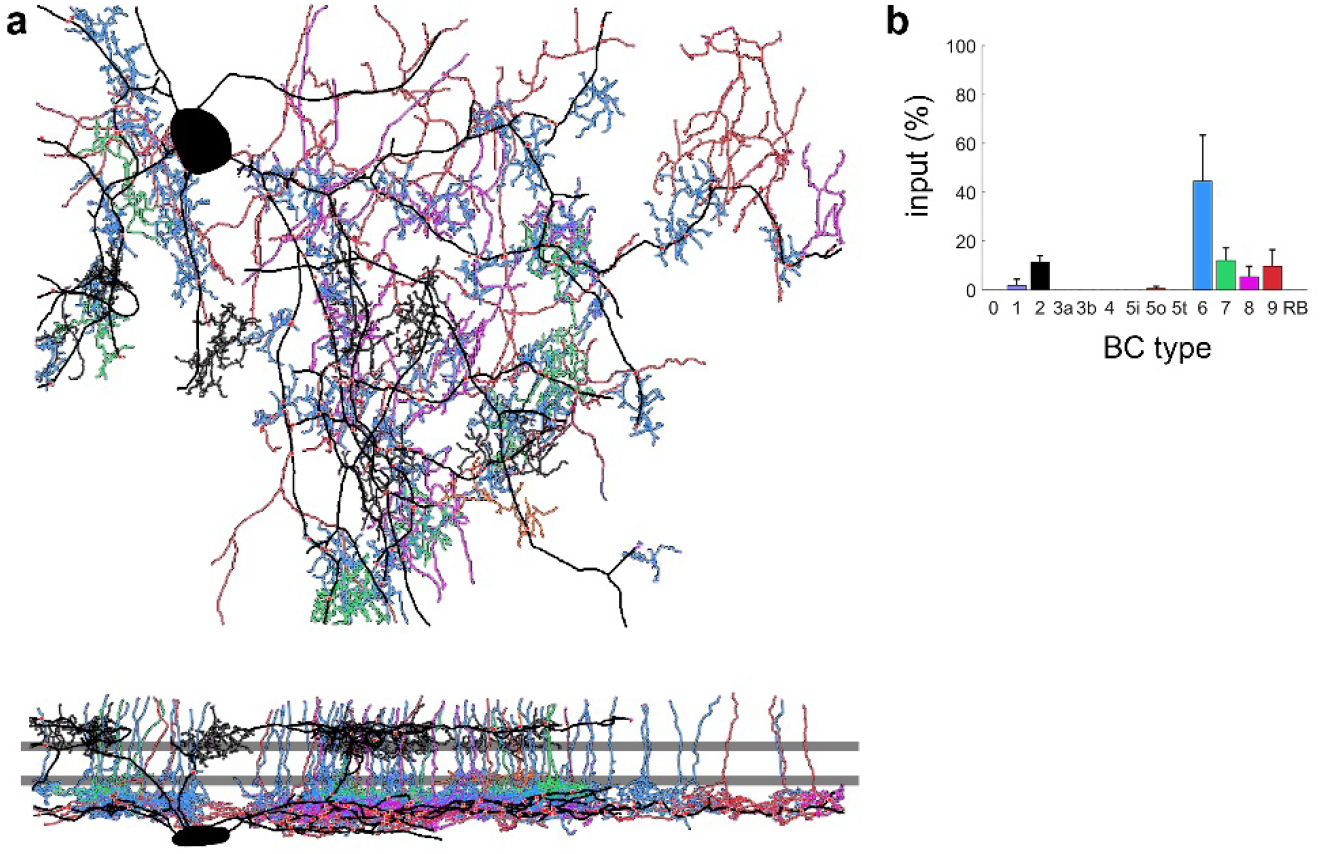
SBEM reconstruction of a bistratified RGC that receives ectopic synaptic input. **a**, SBEM reconstruction of BC input onto a bistratified RGC, as viewed at the plane of the retina and an orthogonal view (bottom). RGC is depicted in black; each BC type is depicted in a unique color. Ribbon synapses are depicted as red circles. Two horizontal gray stripes mark the ON and OFF cholinergic bands. **b**, Percent input of each BC type to the bistratified RGC in (**a**).

**Extended Data Figure 6.**
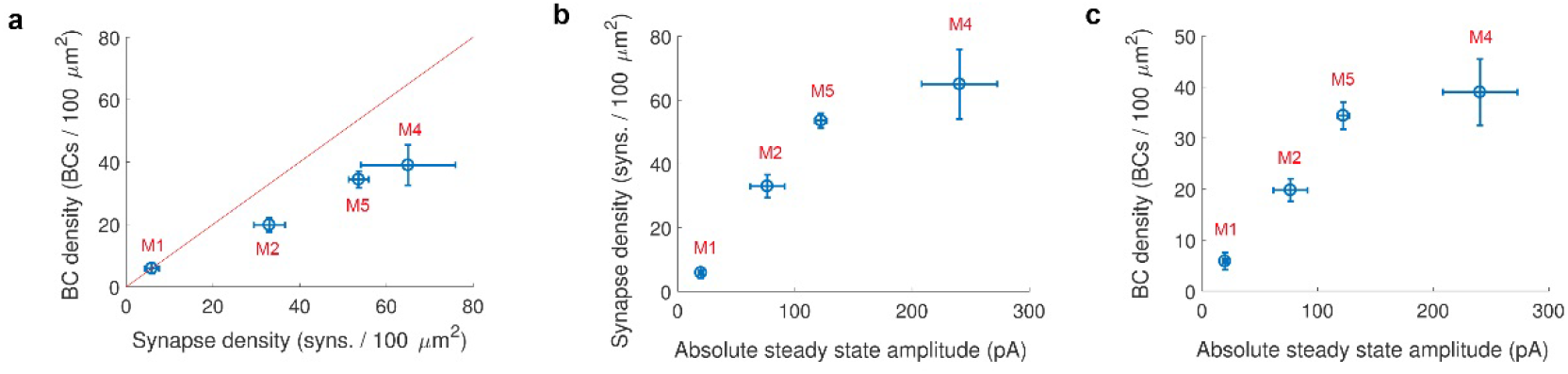
Relationship between amplitude of steady state light-evoked excitatory currents and extent of synaptic input. **a**, Bipolar vs. synapse density (mean ± SEM) across ipRGC types. M1 ipRGCs showed a 1:1 ratio between number of BCs and synapses; data point falls on the identity line (red). However, in all other ipRGC types, the number of synapses was ~1.6 greater than that of BCs; data points fall below the identity line. That is, single BCs made, on average, more than a single synapse onto a given ipRGC. **b**,**c**, Synapse (**b**) and BC (**c**) density qualitatively correlated with the amplitude of steady-state light-evoked excitatory currents in ipRGCs (currents presented in Extended Data Fig. 1).

**Extended Data Figure 7.**
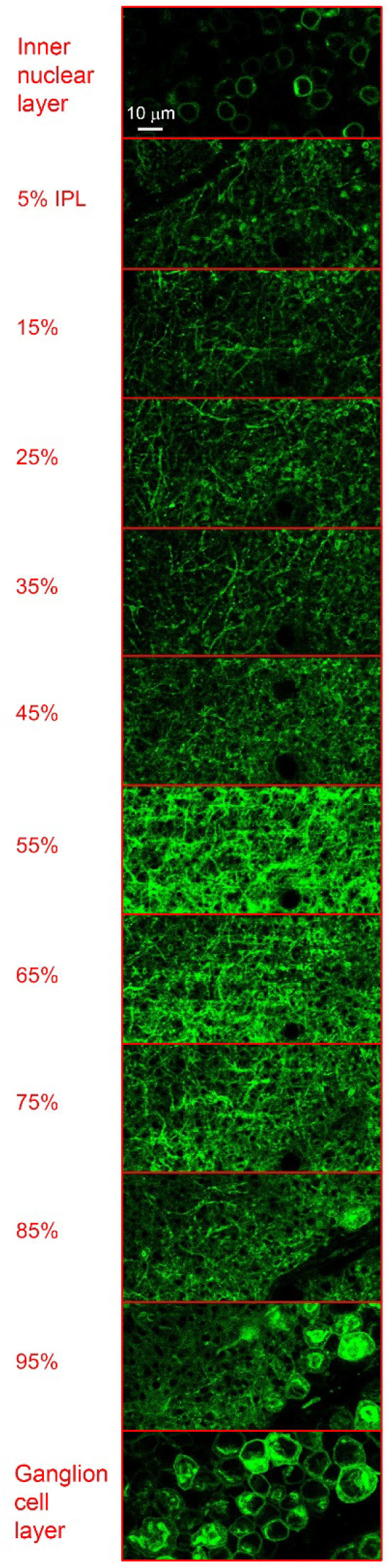
iGluSnFR-expressing dendrites across the IPL. An example two-photon z-stack of the iGluSnFR-expressing dendrites in each examined IPL depth. iGluSnFR expression was highest at the ganglion cell layer and decreased toward the inner nuclear layer. Yet, iGluSnFR expression was high enough to allow reliable measurement of glutamate release at the different IPL depths.

**Extended Data Figure 8.**
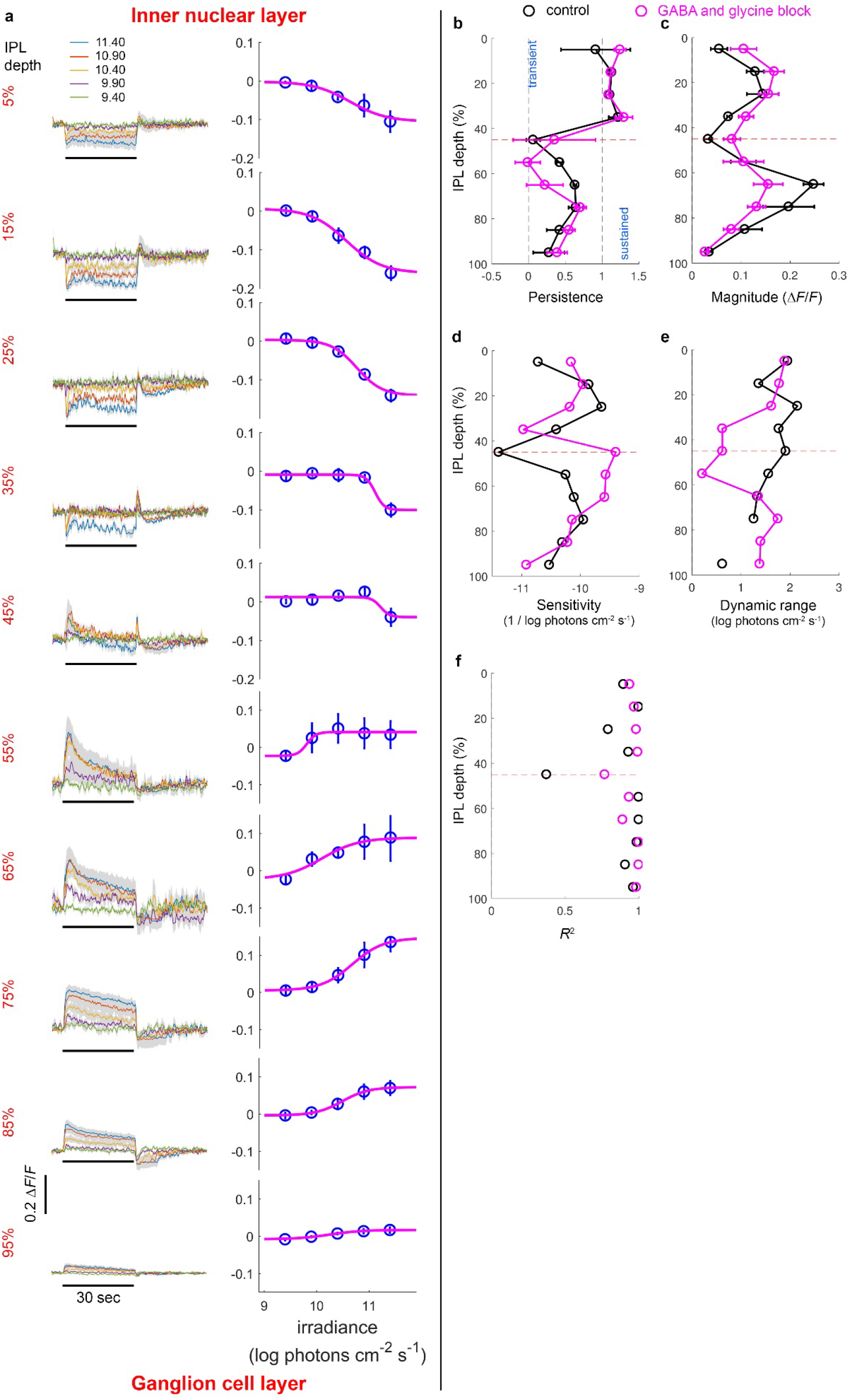
Minimal effect of feedback inhibition on the persistence and capacity for intensity-encoding of BCs’ glutamate release. **a**. Glutamate release from BCs and ACs at 10 different depths across the IPL while blocking GABAergic and glycinergic transmission, from the inner nuclear layer (IPL depth 0%, top) to the ganglion cell layer (IPL depth 100%, bottom). See Fig. 3 for details. Left column shows the glutamate response (Δ*F*/*F*) as a function of time for five light stimulus intensities, for each IPL depth. Right column shows the steady-state glutamate response (Δ*F*/*F*) as a function stimulus intensity, for each IPL depth. BC types that stratify at each IPL depth (whose mean normalized density exceeded 20%) are indicated in black. Blocking GABAergic and glycinergic transmission generally yielded noisier glutamate release traces, especially during the OFF phase of the stimulus for the inner IPL but the ON phase of the stimulus for the outer IPL. That is, such transmission acts as a low-pass filter on the BC glutamate output, removing high-frequency content from the BC signal. Nonetheless, glutamate release still correlated to light intensity both in the inner and outer IPL. Additionally, under blockade of GABAergic and glycinergic transmission, glutamate release in the inner IPL lacked the sharp transient observed under control conditions. Instead, glutamate release gradually decreased with time until light offset, when glutamate release dropped abruptly past the baseline glutamate release. Thus, GABAergic and glycinergic transmission has a role in generating the glutamate transient at light onset. This is reminiscent of the effect of picrotoxin and strychnine on rod light-evoked responses, abolishing a hyperpolarization transient, and instead, generating a depolarization transient at light onset (Szikra et al., 2014). **b**. The persistence index (Per; see Methods) while blocking GABAergic and glycinergic transmission was largely similar to that obtained under control conditions. Per was close to zero throughout the outer IPL (depths 65%-95%), indicating that glutamate release stayed highly sustained also while blocking GABAergic and glycinergic transmission. In the inner IPL, on the other hand, the persistence index at IPL depths of 55% and 65% was higher than under control condition. Thus, at this particular IPL depths, blocking GABAergic and glycinergic transmission made the glutamate release more sustained. **c**. Blocking GABAergic and glycinergic transmission reduced the magnitude of the glutamate signal across the inner IPL, but most noticeably at an IPL depth of 65% (0.15 vs. 0.25 *ΔF*/*F* for treatment and control, respectively). No equivalent reduction in magnitude of the glutamate signal in outer IPL was observed. Otherwise, the magnitude varied across the IPL as it did under control conditions. **d**. Under blockade of GABAergic and glycinergic transmission, sensitivity varied across the IPL roughly the same as under control conditions, but lower sensitivity values were observed in the inner IPL and at the ON-OFF transition depth. **e**. Under control conditions, the dynamic range of glutamate signals varied slightly across the IPL, ranging 1-2 log photons cm^-2^ s^-1^. However, after blocking GABAergic and glycinergic transmission, the dynamic range at the middle of the IPL (depths 35%-55%) was lower by a least 1 log unit than under control conditions. **f**. Blocking GABAergic and glycinergic transmission had little effect on the goodness of fit to the Naka-Rushton function.

**Extended Data Figure 9.**
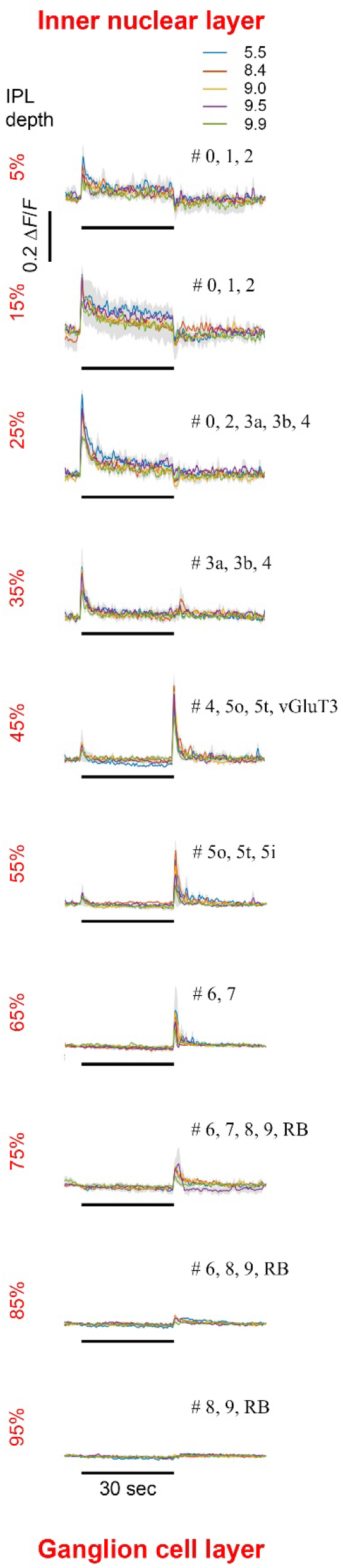
Glutamate release in response to dark spots over a bright background. Glutamate release from BCs and ACs in response to dark spots over a bright background, at 10 different depths across the IPL, from the inner nuclear layer (IPL depth 0%, top) to the ganglion cell layer (IPL depth 100%, bottom). See Fig. 3 for details. For each IPL depth, glutamate release (Δ*F*/*F*) is presented as a function of time for dark spots of five light stimulus intensities (indicated at the top) over a bright background (11.95 log photons cm^-2^ s^-1^). BC types that stratify at each IPL depth (whose mean normalized density exceeded 20%) are indicated in black. Due to inherent technical limitations of our experimental setup, studying glutamate release in response to dark spots (rather than bright spots) was possible only over a limited range of effective stimulus intensities (1.5 log units). This narrow intensity range together with biological and experimental variation did not allow us to test whether glutamate release correlated with stimulus intensity. Nevertheless, the persistence of responses could be quantified accurately. In the outer IPL (5%-35% IPL depth), the glutamate response typically started with a large-amplitude transient, and then gradually rolled off until the termination of the stimulus, upon which glutamate release abruptly dropped to, or at certain IPL depths, below baseline. In contrast, in the inner IPL (55%-95% IPL depth), glutamate release dropped slightly at the onset of the stimulus, and stayed virtually constant until the termination of the stimulus, upon which glutamate release abruptly increased above baseline and spiked. Glutamate release at the ON-OFF transition depth (35%-45% IPL depth) represented summation of glutamate release in both outer and inner IPL.

**Extended Data Figure 10.**
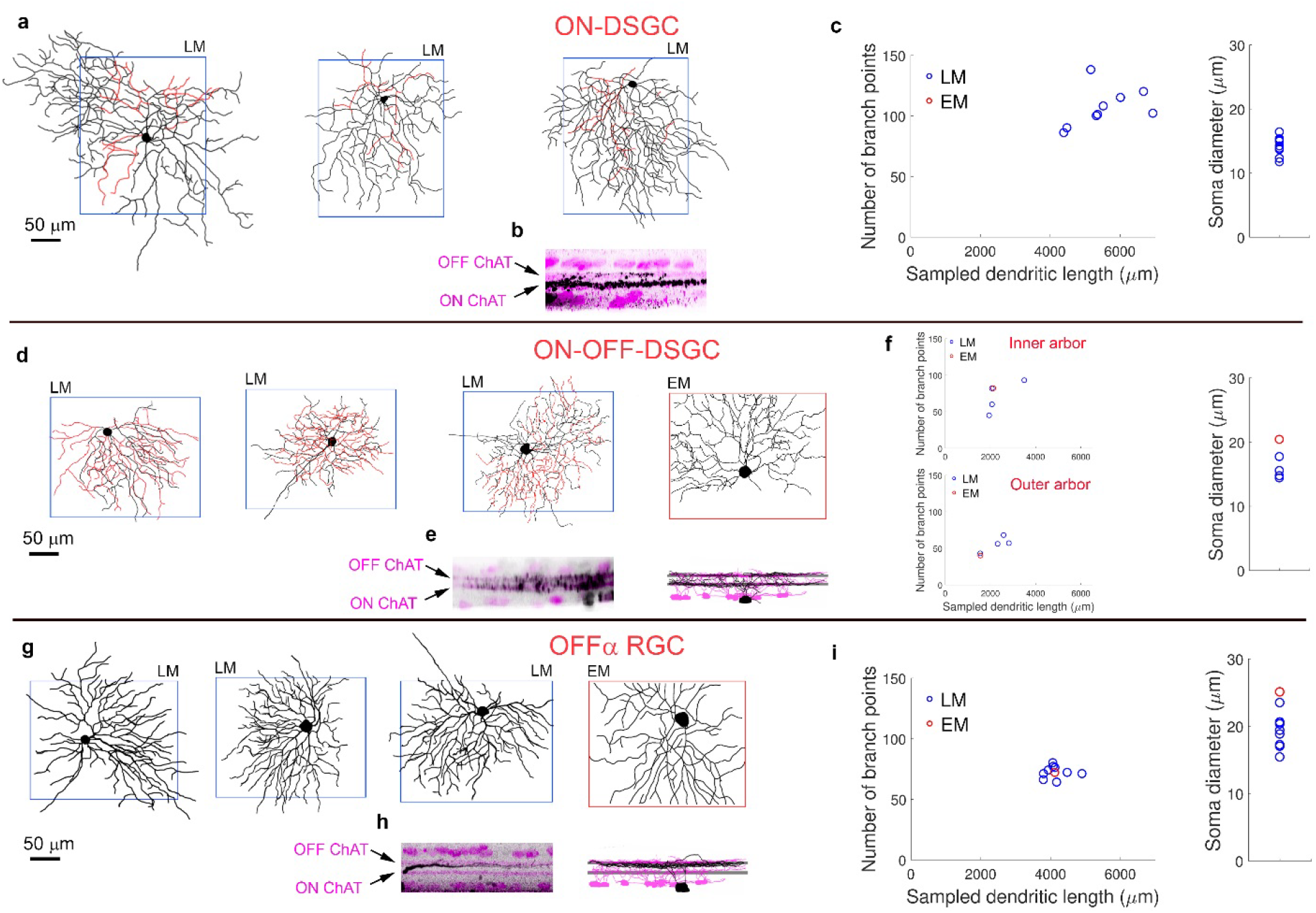
Morphological comparison between SBEM-reconstructed and genetically identified conventional RGCs. **a**. Comparison of selected morphological statistics between SBEM-reconstructed (EM) presumptive ON-DSGC and ON-DSGCs identified using light microscopy (LM) in the Pcdh9-Cre mouse. Three left traces (enclosed by a blue rectangle) represent the *en face* projections of three ON-DSGCs based on depth series (z-stacks) of confocal light micrographs. Fourth trace from the left (enclosed by a red rectangle) represents a presumptive ON-DSGC identified in the SBEM volume. **b**, **left**, Orthogonal projection of a representative LM-reconstructed cell (black) overlaid with anti-ChAT immunostaining (magenta). **b**, **right**, Orthogonal projection of a representative EM-reconstructed cell (black) overlaid with reconstructed ON-OFF DSGCs whose dendritic arbors roughly mark the two ChAT bands formed by the processes of ON and OFF starburst amacrine cells (magenta). The stratification profile of the LM- and EM-reconstructed cells are similar. **c**, **left**. Comparison of the number of branch points as a function of sampled dendritic length for LM-reconstructed (blue) and EM-reconstructed (red) cells. **c**, **right**. Comparison of the soma size between LM-reconstructed (blue) and EM-reconstructed (red) cells. We clipped LM traces using a rectangular frame the size of the SBEM volume (260 μm × 210 μm). Thus, both EM and LM traces sampled an equivalent area of a cell’s dendosomatic profile. See Extended Data Figure 4 for details.. **d**-**f**, The same as **a-c** but for ON-OFF-DSGCs. **g**-**i**, The same as **a-c** but for OFFα RGCs.

